# DNAffinity: A Machine-Learning Approach to Predict DNA Binding Affinities of Transcription Factors

**DOI:** 10.1101/2022.07.26.501522

**Authors:** Sandro Barissi, Alba Sala, Milosz Wieczor, Federica Battistini, Modesto Orozco

## Abstract

We present a physics-based machine learning approach to predict *in vitro* transcription factor binding affinities from structural and mechanical DNA properties directly derived from atomistic molecular dynamics simulations. The method is able to predict affinities obtained with techniques as different as uPBM, gcPBM and HT-SELEX with an excellent performance, much better than existing algorithms. Due to its nature, the method can be extended to epigenetic variants, mismatches, mutations, or any non-coding nucleobases. When complemented with chromatin structure information, our *in vitro* trained method provides also good estimates of *in vivo* binding sites in yeast.

## INTRODUCTION

Proteins are the main regulators of gene expression as they can directly or indirectly inactivate, activate, or enhance the transcription of DNA. Central elements in this regulatory system are transcription factors (TFs): modular proteins that recognize sequences of DNA (typically 6-20 base pair long), helping to recruit RNA polymerases that trigger the subsequent transcription of a nearby gene (1, 2). The binding of TFs during normal cell life is difficult to predict (3, 4) as it is modulated by a myriad of effects, such as the presence of nucleosomes (which in general hinders TF binding) (5, 6), or the formation of clusters that foster cooperativity, general chromatin compaction, or even phase separation (7–9). However, a key requirement for *in vivo* binding is a good binding to the targeted naked DNA.

The recognition of naked DNA by transcription factors is complex and does not follow a common code or a single mechanism (1). Based on the degree of structural distortion that protein induces in DNA, we can distinguish three binding paradigms (10): i) Fischer’s lock and key theory (no distortion on DNA from its canonical B-form); ii) conformational selection (small to medium deformation that aligns with intrinsic deformation patterns of DNA); and iii) induced fit (large deformations of DNA that are unlikely to happen in the absence of the protein). Based on the type of contacts used for DNA recognition, we can distinguish between TFs interacting mainly with the DNA backbone, those establishing hydrogen bond interactions with the nucleobases in either major or minor groove, and finally those disrupting the duplex geometry to generate stacking contacts. In a similar vein, Rohs and coworkers (1, 11, 12) defined two main mechanisms for “TF-DNA reading”: the direct readout, related to the formation of specific interactions of the TF with the nucleobases (typically by means of hydrogen bonds), and the indirect readout, related to the sequence-dependent shape of DNA. Recently, our group extended these ideas by also considering the sequence-dependent energy cost for changing the DNA conformation from the unbound to the bound state (10), a concept that introduces sequence-dependent flexibility as a determinant of sequence-dependent binding.

Most data on the sequence preferences of TFs, i.e. the TFBSs (transcription factor binding sites), rely on high-throughput experimental techniques such as high-throughput SELEX experiments (HT-SELEX; (13–15)), protein binding microarrays (PBM) using either synthetic sequences (uPBM; (16)) or sequences from the genomic context (gcPBM; (17)), or fluidic engines such as HITS-FLIP (18). The final output of these *in vitro* techniques is a list of DNA sequences of different lengths (depending on the experimental method, from 10 in HT-SELEX to 36 in PBM) with an associated estimate of their binding affinity for a given TF. The *in vivo* preferences are typically derived from ChIP-seq approaches (or variants such as ChIP-exo) (19–21), where the chromatin is immunoprecipitated by TF-specific antibody and the retained chromatin is then sequenced. The technique is noisy, as it uses cell populations, and the resolution is poor (200 bp ± 300 in ChIP-seq, and down to 50 bp in the case of ChIP-exo) (20, 21), but it provides direct evidence of the active TFBS under physiological conditions.

Traditional theoretical approaches to predicting TFBS are based on Positional Weight Matrices (PWM) and their associated logos (22, 23). While original versions assume the independence of nucleotide preferences at each position, the newest PWM models can capture some of the sequence interdependence (23, 24), increasing their reliability. However, the limitations of PWM models are well known (25, 26), which has fuelled the development of alternative methods relying on an extended set of descriptive parameters and last-generation learning models. We can broadly classify these new predictive models based on whether they aim to predict *in vivo* or *in vitro* TFBS. Those for *in vitro* TFBS prediction use the nucleotide sequences and a variety of DNA shape descriptors as input parameters (12, 26, 27). Methods for *in vivo* TFBS prediction are typically focused on human genomes, and complement the descriptors of naked DNA (typically sequences) with ones related to chromatin structure and dynamics (RNAseq, DNase, conservation profiles,…). Experimental data have been widely used to train a variety of Machine Learning and Deep Learning methods (28–38) to predict TFBS.

We present here a physics-based ML approach to TF binding affinity prediction *in vitro* that uses the physical properties of DNA directly derived from molecular dynamics (MD) simulations (39–41). These properties consider both the equilibrium geometry and the flexibility, as defined by Olson (42), to describe DNA conformation with sequence-dependence at a base-pair step (bps) resolution. The importance of these conformational properties for the study and prediction of DNA behavior and preferences has been proved in numerous studies (42–48).

Our method uses these DNA physical properties to train a random forest regressor to reproduce uPBM, gcPBM and HT-SELEX data for a large variety of protein families, and yields results that outperform currently available methods. We then use this *in vitro* trained method to explore *in vivo* binding sites of the transcription factor CBF1 in yeast, one of the very few TFs for which we have experimentally available data for both *in vitro* (PB-exo) and *in vivo (*CHIP-exo) binding. When *in vitro* predictions were combined with chromatin structure as determined by nucleosome positioning maps, our method has shown state-of-the-art predictive power in identifying *in vivo* TPBSs.

## MATERIALS AND METHODS

### Datasets for training and testing

There is a variety of data on *in vitro* TF binding preferences, covering a variety of proteins and measuring techniques. <u>HT-SELEX data</u> were taken from 2 different studies (European Nucleotide Archive ENA PRJEB14744 and PRJEB29730) and processed using the associated package (https://bioconductor.org/packages/release/bioc/html/SELEX.html). The combined databases contain information on 600 TFs from more than 30 different protein families. TFs with no k-mer (10 bases) with at least 100 counts in 0-th order SELEX cycle were removed from the training set as they have no clear binding motif (as discussed in (13)). As reported in the literature (13), training and testing were done using data from the penultimate SELEX cycle provided in the databases. <u>uPBM data</u> were taken from the DREAM5 challenge containing information on binding preferences of 35-mer oligos for 66 mouse TFs (49). The 50 oligos with the highest affinity to a given TF were used to define a PWM (50), which was then used to align the sequences and derive the most probable binding site (a shorter k-mer, typically 12-mer). Finally, for gcPBM data, sequences already aligned around the putative binding site placed at the center of 36mer genomic were used; more specifically, sequences for the TF dimers Mad1/Max (‘Mad’), Max/Max (‘Max’), c-Myc/Max (‘Myc’) and CFB1 (Gene Expression Omnibus accession numbers GSE59845 and GSE44604 respectively) (17, 51–53). Possible multiple binding sites were removed using a previously published protocol (26).

*In vivo* binding data for CBF1 were taken from ChIP-exo maps of *Saccharomyces Cerevisiae* (GSE44604 and GSE147927) (17, 52, 53) and were used to perform a proof of concept of the ability of the in vitro CFB1 binding predictor to detect in vivo binding sites. Data on chromatin used to discuss the differences between *in vitro* and *in vivo* binding profiles were taken from previously published nucleosome maps in the same cellular model (5, 54).

All ID and references to the datasets used for training and testing are summarized in Supplementary Table S1.

### Feature Classes

For the training of the machine learning (ML) algorithm, different classes of features were used:

- ***Sequence composition: Presence*.** For each k-mer, a vector of counts for the 256 possible tetramers that show up in a given k-mer was calculated, using a sliding window of length 4 and simply adding occurrences. For instance, the 6-mer ‘AAAAAT’ would have two counts for the tetramer ‘AAAA’, one for ‘AAAT’ and none for the remaining 254 tetramers.
- ***Indirect readout: Base pair parameters*.** The base pair parameters (equilibrium values and the diagonal components of the stiffness constant matrix, called AVG and DIAG respectively, (37–39)) for each individual base pair step movement (translational: shift, slide and rise, and rotational: tilt, roll and twist) were considered. The values were retrieved from a dataset that covers all the unique base pair steps in all the possible tetranucleotide environments from microsecond-long molecular dynamics simulations (10, 41). All data used to characterize tetramers are available in our BigNASim database (55).
- ***Direct readout***. The electrostatic patterns of each base (hbond acceptor/donor or hydrophobic) were considered using the scheme below (see Supplementary Figure S1) (12). Our method assigns integers (−1, 0, +1) to acceptors, hydrophobic sites and donors respectively, and for each overlapping tetramer along the DNA sequence sums the relative values along columns. For the minor groove, since the flanking sites are always −1, they are omitted and we keep only the middle value. In total, for each tetramer the electrostatics are explained by 5 values, for example, for ‘AACT’ [−1,+1,−1,0,0] + [−1,+1,−1,0,0] +[0,+ 1,−1,−1,+1] + [0,−1,+1,−1,0] = [−2,2,−2,−2,1].

All the references to the features used are summarized in Supplementary Table S1.

### Machine Learning Training

Features described above were used as descriptors, and they were attributed to each overlapping tetramer in the k-mer sequence under study (see Supplementary Figure S2). Experimental affinities in databases were used as labels for a Random Forest regressor (56). A train/test ratio of 80/20 on the experimental data (HT-SELEX, uPBM and gcPBM) (see Scheme) was used to reduce overtraining artifacts. The R^2^ regression scoring function, Pearson correlation (r) and MSE (mean squared error; see Supplementary Methods for details) between the predicted and experimental affinity (56) was used to validate the accuracy of the model (Figure 1). For the training process, we randomly selected 80% of the data, and we performed bootstrap to avoid biases. This method allowed us to perform the training multiple times even if the chosen training and testing sets could have contained repeated entries. For validation of the method, we also tested the choice of the training set using cross-validation, which performs the simulation K times dividing the data into K partitions and using each time one different partition as a test set. Changing the algorithm to K-fold cross-validation (K=10; 90/10 randomly chosen), we obtained very similar results as previously (see Supplementary Table S2).

**Figure 1.**
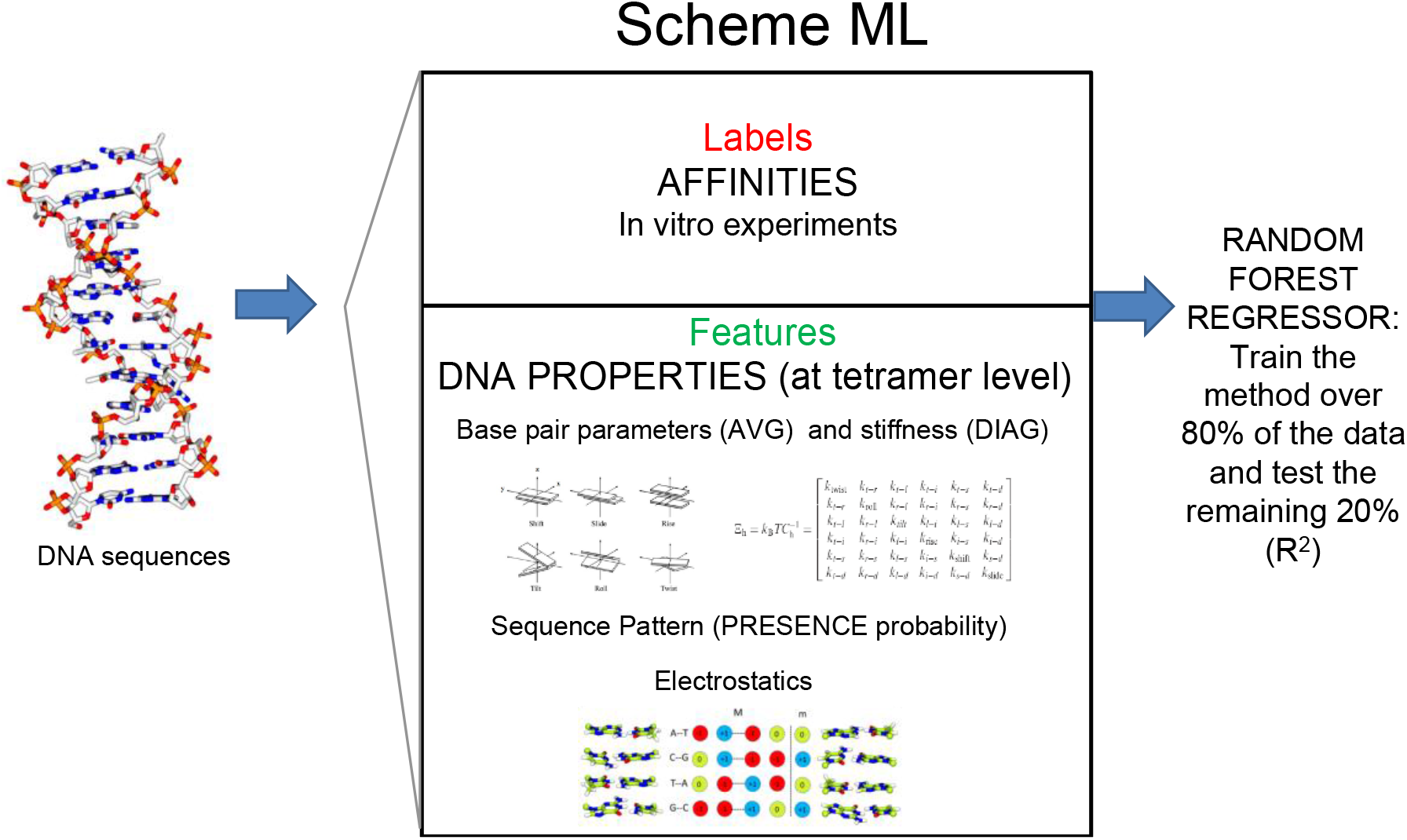
General scheme of the ML training strategy

To improve the accuracy and efficiency of the training process and to avoid training artifacts, we applied several data pre-processing steps:

**Undersampling.** The uPBM experimental datasets are usually very noisy with an overpopulation of k-mers with a low affinity value. To obtain a more balanced set, we applied different undersampling approaches for each dataset type (details and an example in Supplementary Methods and and an example in Supplementary Figure S3)..
**Weighting.** For the uPBM dataset, because of the lack of high-affinity sequences, we assigned uniform higher weights (10) to sequences matching the PWM data for each TF being considered. We also calculated the importance of each class feature in the regressor (56).

The purpose of undersampling and weighting (or oversampling of high affinity values) is to remove noisy data by stratifying the affinity profile and picking only a few samples in each stratum.

### HT-SELEX data quality assessment

For HT-SELEX data, we considered the target affinity values as those reported in the cycles. In principle, binding affinities of the different k-mers should grow exponentially at each cycling step until saturation, and any deviation from this behaviour signals inconsistency of the data. Thus, to guarantee the quality of the experimental data, we performed a quality check with a Support Vector Machine (SVM) discriminant using the correlation between selected counts across different cycles as descriptors (see Supplementary Methods).

### Genomic testing

For the *in vivo* validation on the yeast genome, we generated discrete maps using the relative score obtained from the ML training for each possible TFBS and comparing the predictions with *in vitro* (PB-exo) (17) and *in vivo* maps (ChIP-exo) (17, 52, 53). We define:

- True positive (TP) when the experimental peaks (both PB-exo and ChIP-exo) overlap with our prediction.
- False positive (FP) when TFBSs predicted by our model do not correspond to any experimental peak. We also applied this category if our prediction corresponds to just one of the experimental dataset (either PB-exo or ChIP-exo)
- Nucleosome occupied locations (Nuc) when comparing our prediction with nucleosome maps one TFBS predicted by our model overlaps with a nucleosome.
- False negative (FN) when experimental TFBS peaks have not been predicted by our model.
- True negative (FN) when both experiments and predictive model agree that there is not a TFBS.

The program and the full database of feature parameters are available in the GitHub repository: https://github.com/Jalbiti/DNAffinity.

## RESULTS AND DISCUSSION

For each experimental dataset (gcPBM, uPBM and HT-SELEX), we trained our machine learning regressor to predict experimental binding affinities using three classes of features informative of the three DNA-protein recognition modes: sequence, direct and indirect readout. After training on 80% of the data, we calculated the determination coefficient (R^2^) between our predictions and the experimental values (see Methods) on the remaining 20% of the data. With that, we found that our model is able to reproduce gcPBM data with astonishing quality, as shown in an average R^2^ of 0.93±0.02 (see Figure 2).

**Figure 2.**
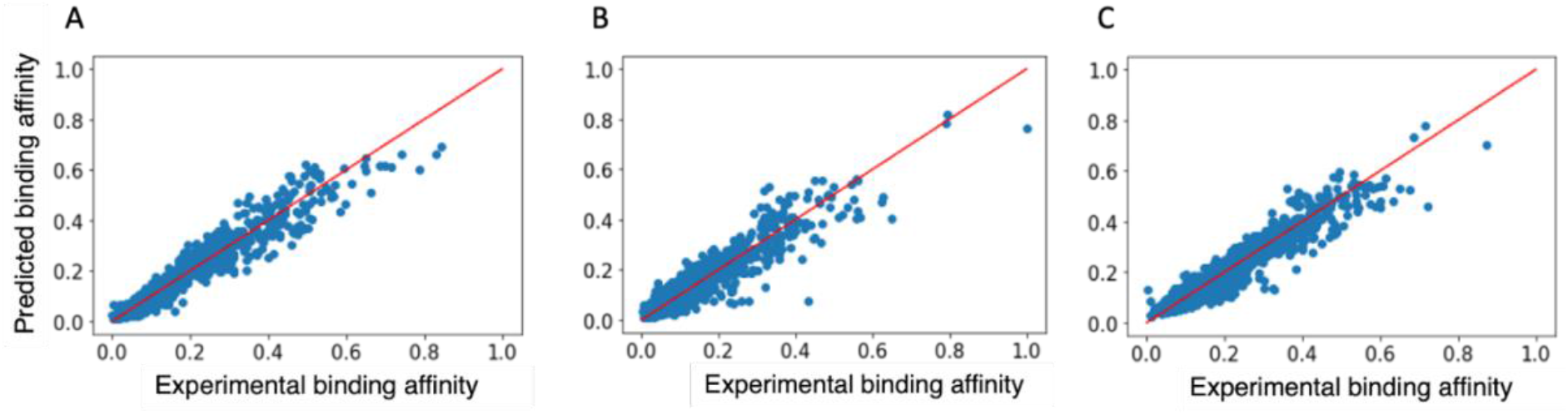
Correlation between the predicted and experimental TF binding affinity for the three cases under study: A) Mad1/Max (‘Mad’), B) Max/Max (‘Max’) and C) c-Myc/Max (‘Myc’). Correlation with experimental affinities for the transcription factors: MAD1 (left panel): R2=0.951, MYC (central panel): R^2^=0.905, MAX (right panel): R^2^=0.922.

Using the <u>uPBM</u> data as reference (see Methods), we could predict affinities with an average determination coefficient R^2^ 0.69±0.17 (Figure 3).

**Figure 3.**
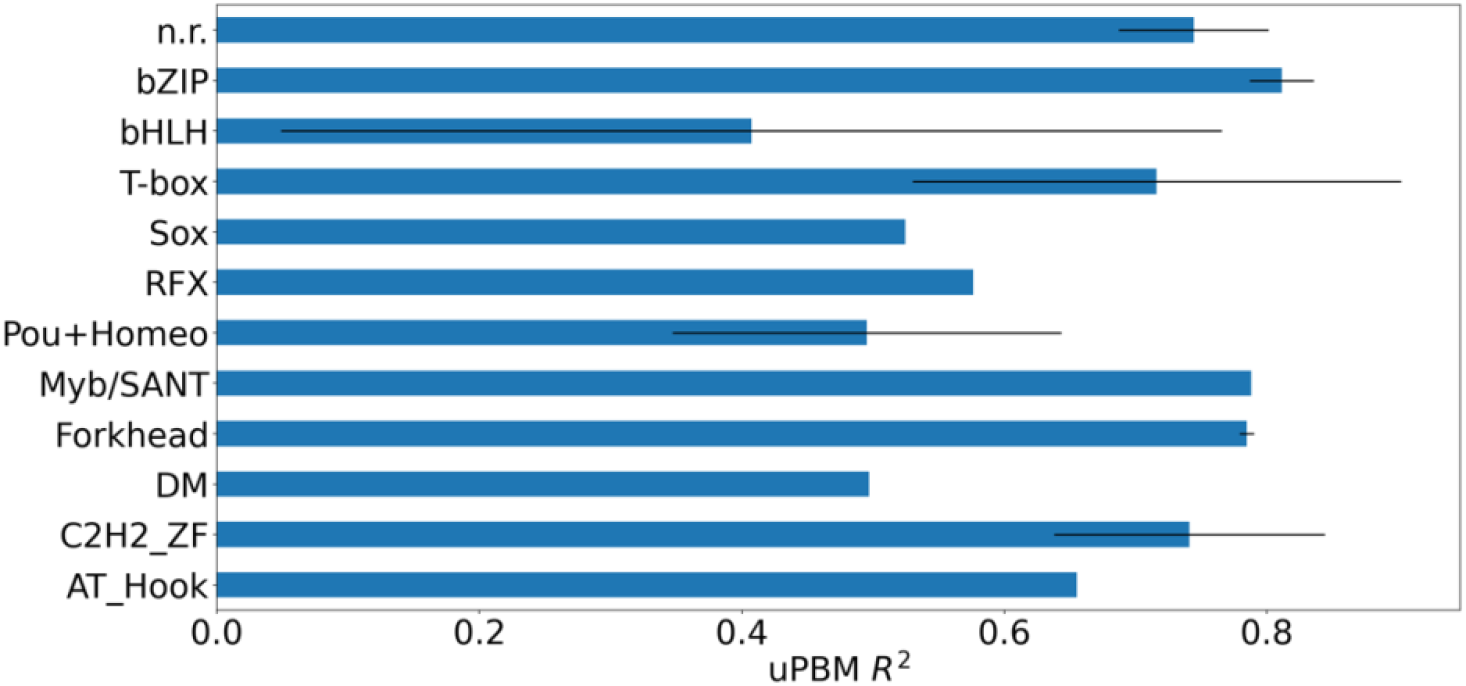
Determination coefficient (R^2^) between predicted and experimental data for different protein families using uPBM data.

We then applied our ML predictor to two datasets from HT-SELEX (see Methods and Supplementary Table S1): the first using the results based on 5-cycles of HT-SELEX experiments, and the second on 7-cycles. In the first case, we achieved an R^2^ of 0.63±0.19, and in the second 0.71±0.21, which yielded a total average of 0.66±0.19. Supplementary Figures S4-S7 detail the results for each HT-SELEX experiment and each transcription factor. In a further step, we used SVM to remove those cases displaying inconsistency between the enrichments at different HT-SELEX cycles (see Methods and Supp. Methods) as they are suspicious cases whose inclusion can bias the training and testing. The improvement obtained after SVM-filtering of data is clear, as seen from the average R^2^ of 0.70±0.14 and the dramatic reduction of cases with low R^2^ (see Figure 4).

**Figure 4.**
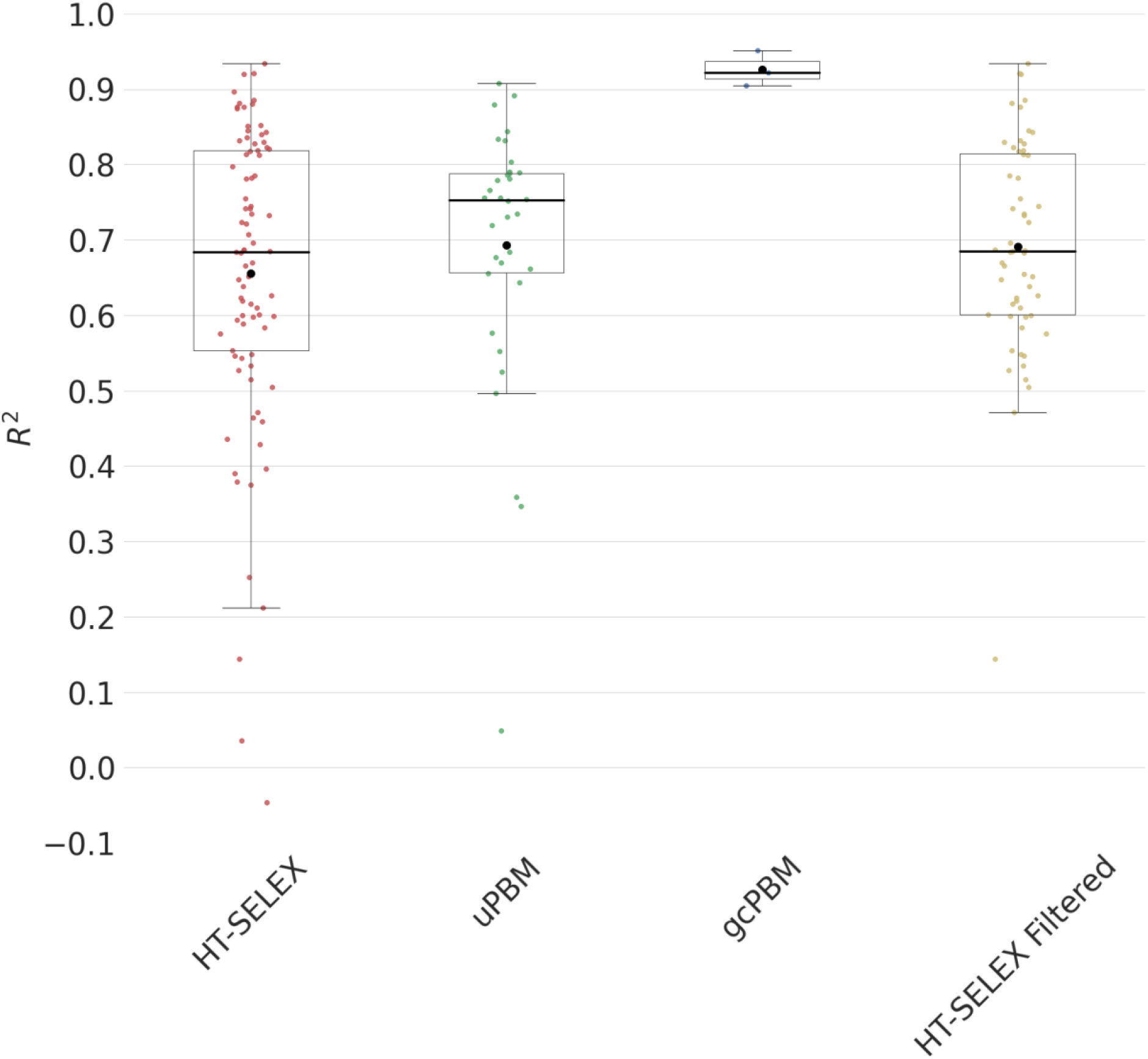
Determination coefficient (R^2^) between predicted and experimental affinities for different transcription factors (black circle marks the mean, black bar marks the median value). Data used were: HT-SELEX on the entire dataset, uPBM, gcPBM, and HT-SELEX on the filtered dataset (after applying our reliability filter that uses an SVM classifier on the raw data; see Methods and Supplementary Methods).

In summary, DNAffinity is able to accurately predict relative binding affinities of transcription factors with a common set of descriptors, irrespectively of the source of experimental approach used to determine the binding (see Figure 4). The logo plots of the most favorable TFBSs, calculated using the top 100 predicted sequences, for each TF studied are presented in Supplementary Table S3.

### DNAffinity’s performance compared to existing ML predictors

We compared our predictor with a larger variety of previously published methods that combine different learning approaches and include DNA sequence and DNA shape features (27, 35, 36, 49) (Figure 5 and Supplementary Figures S8-S9). In all cases comparisons are done using the same number of cases (TFs) and the same dataset as used by the original developers of the methods.

**Figure 5.**
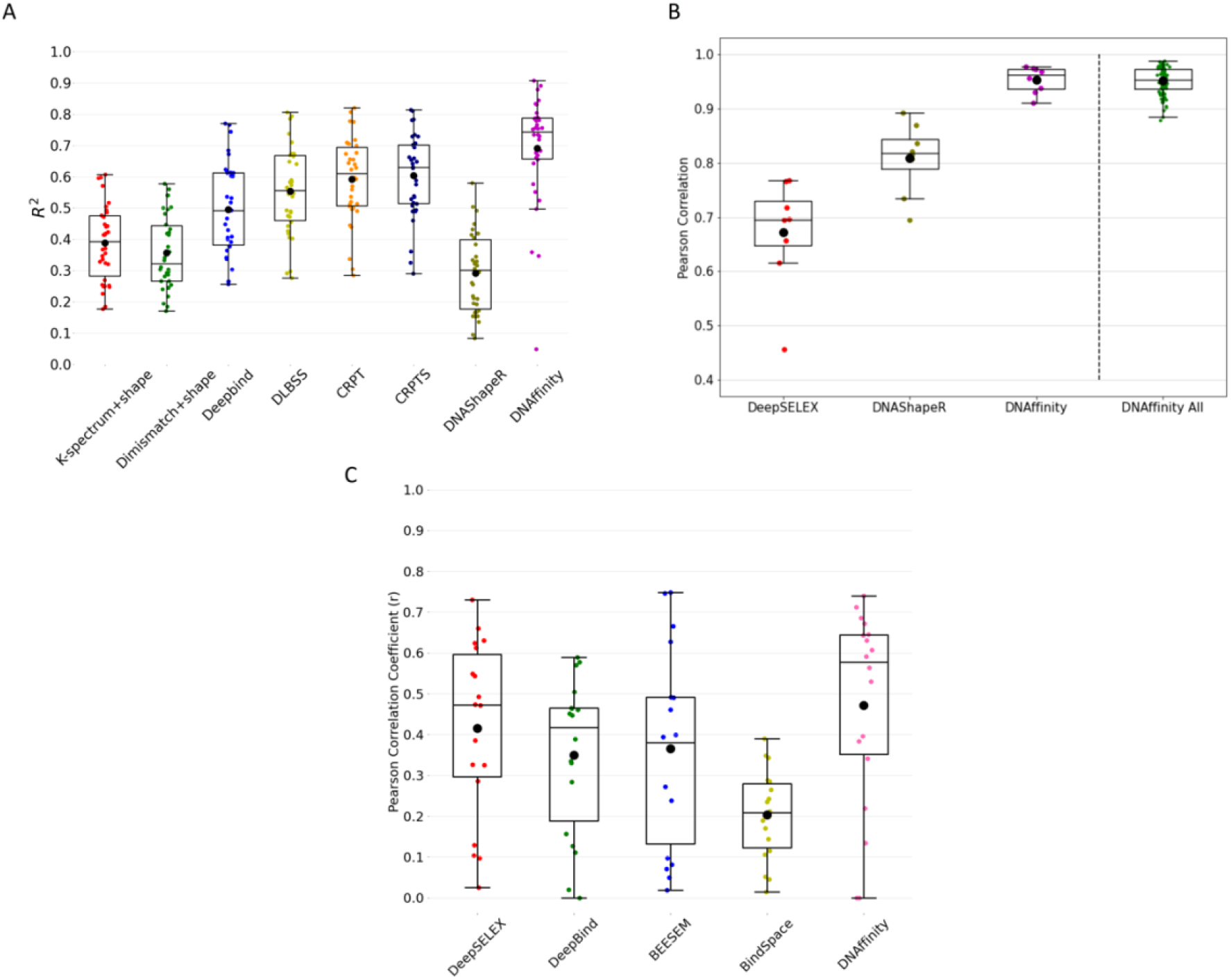
Comparison between our predictions and previously reported ones. A) Determination coefficient (R^2^) between predicted and experimental data for different protein families using uPBM data, data retrieved from (27) and DNAShapeR (33). B) Pearson correlation using in common HT-SELEX data obtained in the literature by previously developed methods, DeepSELEX and DNAShapeR (33, 38) and ours (DNAffinity) respectively, values obtained using all the HT-SELEX data available using our method (DNAffinity all). C) Pearson Correlation between predicted and experimental data for different protein families using HT-SELEX for training and testing on uPBM data, using our method (DNAffinity) and previously developed methods: DeepSELEX (38), DeepBind (36), BEESEM (58) and BindSpace (59). The data reported were taken from (38). Black circle: mean, black bar: median.

For gcPBM our performance (Figure 2) is nearly identical to that obtained with the shape-augmented ML predictor (32), but this outstanding performance should be taken with caution as there are only 3 transcription factors for which gcPBM information is available. Here, the probes are very well aligned based on the central TFBS and the sequences used do not show high variability, making the problem relatively easy to solve.

A more reliable and challenging benchmark can be obtained using uPBM and HT-SELEX data as: i) there are more datasets available and ii) there is a larger variety of trained models to benchmark against.

In order to evaluate the performance of DNAffinity, we compared its predictive power on 66 uPBM datasets and compared our R^2^ with the ones obtained using the same dataset by: CRPTS/CRPT (a hybrid convolutional recurrent neural network (CNN/RNN) architecture that combines DNA sequence and DNA shape features) (27); Deepbind (a CNN model primarly based on DNA sequences) (36); two kernel-based methods (spectrum + shape kernel, di-mismatch + shape kernel) (57); a deep learning-based DLBSS (37); and a shape-based ML regressor DNAShapeR (33). The data used for the comparison were previously reported (27). In Figure 5A and Supplementary Figure S8 we show the results of the comparison. The improvement with respect to all other predictive algorithms is very clear: our algorithm has a stronger predictive power when compared to methods based on shape and neural network, probably thanks to the properties considered and the pre-processing of the 36mers.

For HT-SELEX data, we compared our results with a shape-based ML regressor, DNAShapeR, and DeepSELEX (38) trained and tested on the same TFs (Figure 5B). Running DNAShapeR, we used their refined set of sequences (M-word, https://rohslab.usc.edu/MSB2017/) and their latest parameters. The comparison shows how our method could better predict the TFBS affinities, and that the results we get for those selected protein is in the range of our prediction using all the cases (DNAffinity All in Figure 5B) available for HT-SELEX (see Supplementary Table S1). We tested the rigorousness of our method calculating the MSE (see Supplementary Methods) and comparing it to the other algorithms that had the best (newest) and the worst performance compared to ours: CRPT (27) and DNAShapeR (33) for uPBM data and both DNAShapeR (33) and DeepSELEX (38) for HT-SELEX data (see Supplementary Table 4); confirming that also using this metric the results obtained by our algorithm are consistent.

Finally, many previously developed methods verified the transferability of their predictive algorithm by first training with one dataset (HT-SELEX) and subsequently testing on another dataset (uPBM). These two experimental techniques differ in the variety and number of sequences that can be studied, and on the length of the different probes: HT-SELEX considers a large amount of different short sequences with one possible binding site, while uPBM has fewer and longer probes with multiple candidate binding sites, and mainly low affinity. Consequently we also compared the ability of our method to inter-cross between datasets. The results obtained (Figure 5C) show that when comparing our results to previously published methods, including neural network and deep learning algorithms (data taken from (38)), DNAffinity outperforms all of them. We obtained an average Pearson correlation of 0.47 vs. 0.41 (DeepSELEX) (38), 0.35 (DeepBind) (36), 0.36 (BEESEM)(58) and 0.20 (BindSpace) (59).

### Importance of the features

Contrary to our original expectations, we found that the impact of the different features on the predictive power of the method depends dramatically on the type of experimental data used for training (Figure 6).

**Figure 6.**
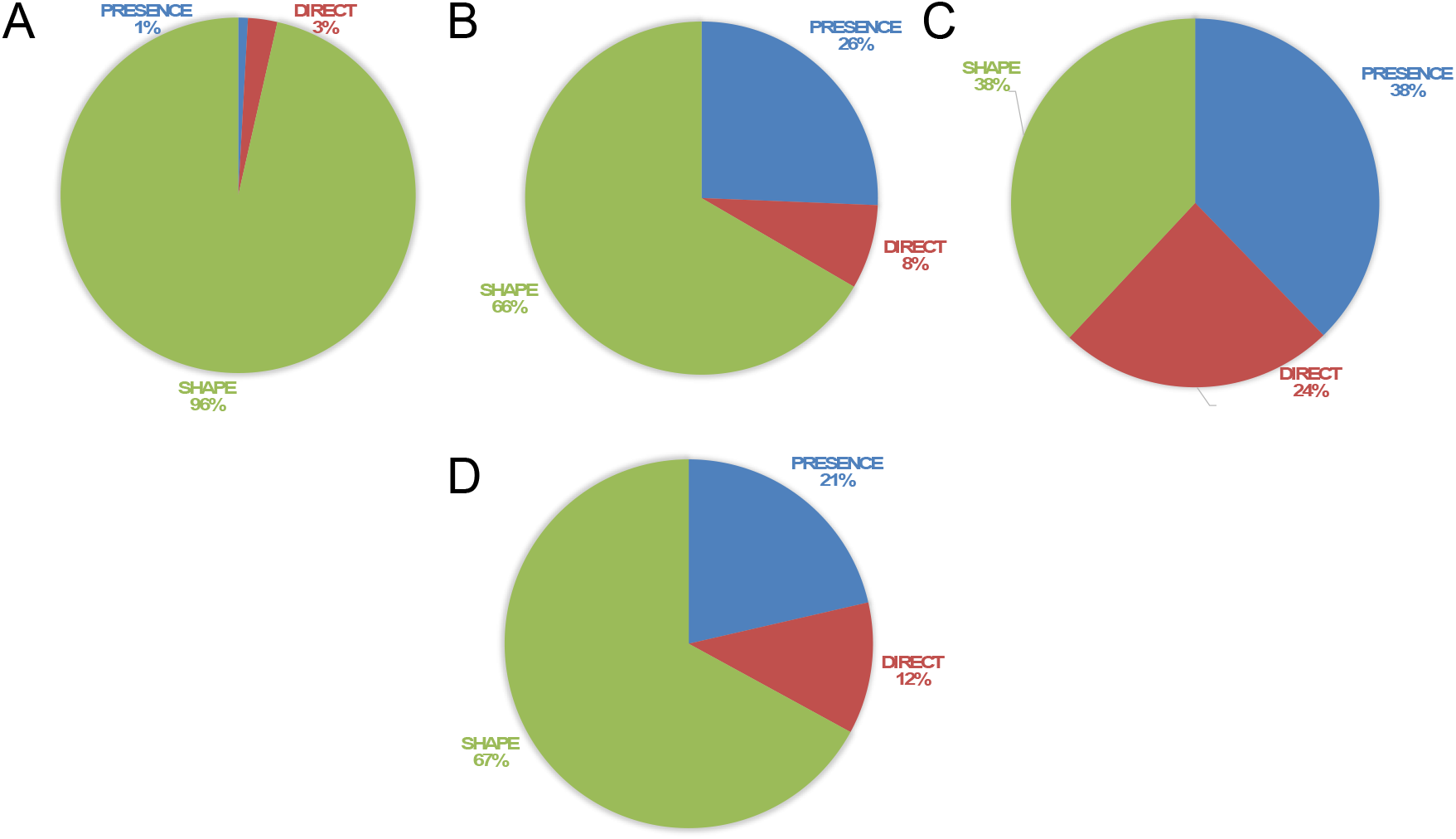
Relative importance (%) of each feature class in the prediction. A-C) Results regarding prediction of gcPBM, uPBM and HT-SELEX data respectively. D) Average of the relative importance (%) considering the prediction on all the datasets (A-C).

For gcPBM data, characterized by a very low sequence variability and a very well defined TFBS, the physical properties of the tetramers have a higher importance, probably because they can differentiate between otherwise very similar sequences.

For uPBM, we trimmed the sequence based on MEME suite result, so the variability of the sequence diminishes. For this reason, in part like in case of gcPBM, the motifs present a common pattern and shape seems to be the best class of feature capable of accentuating their differences (see the contribution of different feature classes in Supplementary Figures S10 and S11). However, because the sequence variability is broader than in gcPBM, also sequence and electrostatics features seem to gain importance.

On the contrary, for HT-SELEX, where a wide range of sequences is reported, the predictive power is equally divided across the three feature classes (Figure 6). This can be explained considering that the method explores a larger range of sequences, including these that are not physiologically accessible. Also, in this case we studied the importance of every class of feature for the prediction and detected that all of them contribute to the prediction (Supplementary Figures S10 and S11). Interestingly, the electrostatic descriptor gained importance when using HT-SELEX data compared to uPBM. The addition of this new “direct-reading” feature to the prediction scheme introduces a new dimensionality in our method to discriminate among largely variable sequences. The HT-SELEX dataset seems pushing the predictive power of the models to their limits.

Above all, by analyzing the effect of each feature on the different protein families it could be possible detect the ones that are mostly affected by indirect-readout (shape and force constants) or direct-readout (sequence and electrostatics) descriptors (see Figure S11). Our results also raise concerns about attempts to train ML methods with a narrow set of sequences, and give us confidence that DNAffinity provides results of very similar quality when reference data come from uPBM or HT-SELEX with a common set of features.

#### In vivo testing

We finally applied our method to simultaneously predict *in vitro* and *in vivo* datasets describing the TFBS for the protein CBF1 (one case for which both *in vitro* and *in vivo* data are available, see Methods). After training our regressor using gcPBM data (R^2^=0.80), we applied our model to predict PB-exo and ChIP-exo peaks along the yeast genome. We considered the consensus exo peaks, because being in common they are independent on the experimental technique/conditions. Each method (ChIP and PB) has some intrinsic noise due to non-specific or spurious interactions and that using consensus peaks ensures that each signal is genuine. To account for the impact of chromatin structure, we include accurate nucleosome maps collected for yeast in the G1 phase (5, 54). Quite encouragingly, we were able to predict almost all consensus exo peaks, defined as locations where PB-exo and ChIP-exo signals coincide (Figure 7). Our true positive (TP) rate (TP/total number of exo experiments peaks) was over 94% (TP case example in Supplementary Figure S12A), meaning that only <6% of the consensus exo peaks are not detected by our method (see Figure 7). Although these TPs entail only 14% of all predictions of our model, a vast majority of the theoretically false positive are at locations occupied by nucleosomes (see Figure 7 and example in Supplementary Figure S12B). As those chromatin sites would not have been accessible for the binding of a transcription factor (occupied by nucleosomes, Nuc in Figure 7B), they correspond to cases where intrinsic (*in vitro}* binding can be favourable, but chromatin structure precludes effective *in vivo* binding. Thus, when nucleosome maps are included as descriptors, the resulting false positive (FP) rate is just 11%. In fact, of the 146 “bona fide” FPs, 37 (FP2 in Figure 7) correspond to sequences that matches one exo signal (PB-exo or CHIP-exo) and 15 have evidence of activity based on polymerase maps (fourth column in the classification scheme Figure 7, FP case examples in Supplementary Figure S12C and D). It means that the real FP rate can be as low as 7% (FP1 and third column scheme in Figure 7). Due to the lack of simultaneous *in vivo* and *in vitro* binding data, it is difficult to generalize our conclusions; we consider here nucleosome occupancy as a proxy for chromatin structure, but there are many other means by which cells can hide regions that would otherwise be bound by transcription factors. Even though we will never be able to make a prediction taking into consideration all the possible variables to *in vivo* TF binding, we transferred the *in silico* prediction to *in vivo* conditions. We think it is important to determine how the intrinsic sequence-dependent binding properties *in vitro* are affected by chromatin accessibility. Our results may suggest that *in vivo* binding may be understood as in vitro binding corrected by high-resolution (nucleosome-scale) chromatin structure.

**Figure 7.**
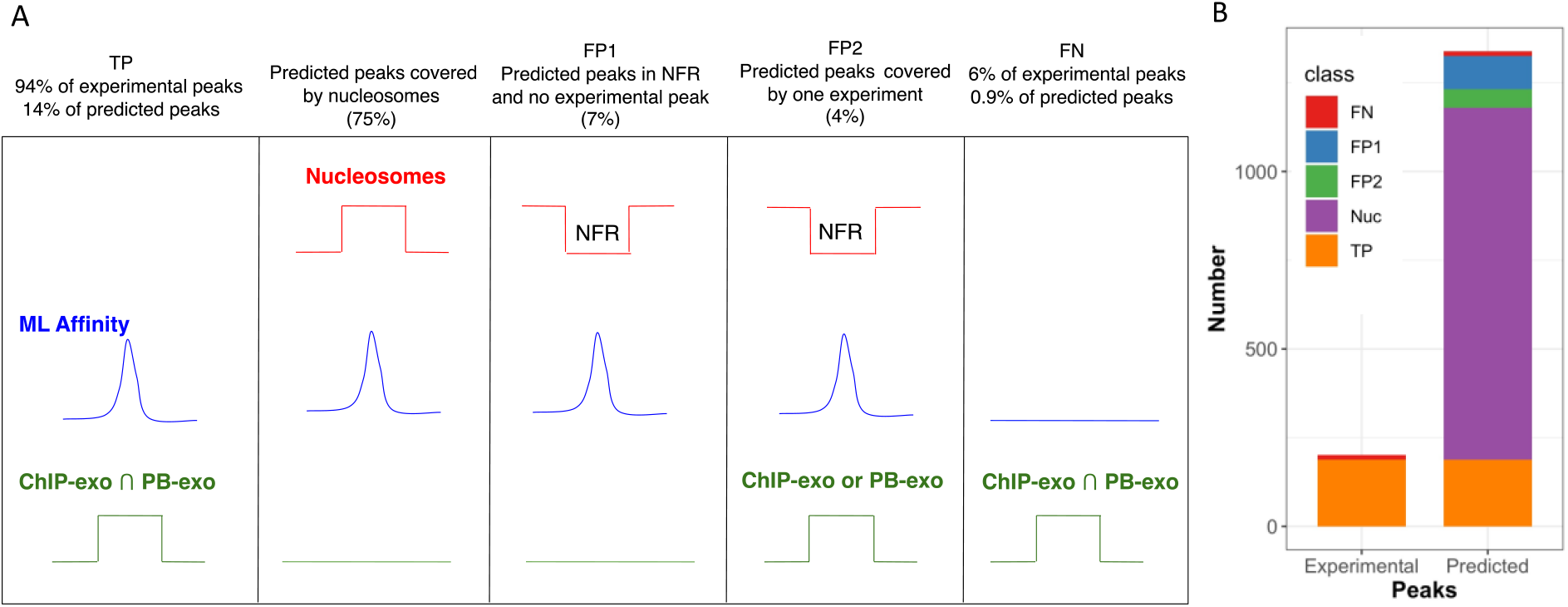
A) Scheme representing the prediction scores along the yeast genome. B) Statistics of our prediction along the yeast genome. On the right, distribution over all the predicted peaks: 188/1326 true positive (TP, orange); 13/1326 false negative (FN, red); 992/1326 in locations occupied by nucleosomes (in purple); the false positive (FP) cases are divided into FP1 that correspond to predicted peaks at nucleosome free region (NFR) that do not correspond to any experimental peak ((94/1326, in blue) and FP2 (52/1326, green) that correspond to sequences that either match one exo signal (PB-exo or CHIP-exo) or have evidence of activity based on polymerase maps. On the right, distribution over all the consensus experimental peaks (PB-exo and CHIP-exo): 188/201 true positive (TP, orange); 13/201 false negative (FN, red).

## CONCLUSIONS

Prediction of transcription factor binding sites is the next grand challenges in genomic research. Development of efficient predictive algorithm requires solving a series of intrinsic problems: on the one hand, the concept of transcription factor binding site is not uniquely defined, as it deeply depends on the intrinsically noisy and low-resolution experimental technique used to detect it, making it impossible to create a universal predictor. On the other, transcription factors use a repertoire of mechanisms for selecting target DNA sequences, and the most informative parameters describing these mechanisms largely depend on the sequence variability explored by the experiment. The complexity of the problem increases even more if *in vitro* predictions are tried to be extrapolated to *in vivo* settings, where other factors besides intrinsic transcription factor affinity play a role.

Our predictive model (DNAffinity) is based on a simple machine learning algorithm trained on *ab initio* parameters derived from first-principle molecular dynamics simulations. One of the advantages of using theoretically derived descriptors is that they can be in principle obtained for any non-coding DNA, including epigenetic variants or lesions. Despite the “ab initio” nature of the descriptors and the simplicity of the training, the method provides excellent results, outperforming all available competitors when predicting *in vitro* transcription factor binding sites irrespective of the experiment used for validation. Very encouragingly, DNAffinity trained on *in vitro* data showed an excellent ability to detect the binding sites of the same transcription factor *in vivo*. Thus, even though DNAffinity predicts many potential binding sites where no experimental evidence of *in vivo* binding exists, a grand majority of these seemingly false positives are trivially explained by chromatin structure and nucleosome occupancy. When combining DNAffinity and nucleosome maps, our method was able to locate *in vivo* TFBS with a high accuracy.

## CONFLICT OF INTEREST

None declared.

## ACKNOWLEDGEMENTS

This research has received funding from the Center of Excellence for HPC H2020 European Commision. “BioExcel-2. Centre of Excellence for Computational Biomolecular Research” (823830), the Spanish Ministry of Science (RTI2018-096704-B-100) and the Instituto de Salud Carlos III–Instituto Nacional de Bioinformatica (ISCIII PT 17/0009/0007 co-funded by the Fondo Europeo de Desarrollo Regional). This project is co-funded by the European Regional Development Fund under the framework of the ERFD Operative Programme for Catalunya, the Catalan Government AGAUR (SGR2017-134). The IRB Barcelona is the recipient of a Severo Ochoa Award of Excellence from the MINECO. Modesto Orozco is an ICREA Academy scholar. SB is a M4L student.

## Notes

https://github.com/Jalbiti/DNAffinity

### Competing Interest Statement

The authors have declared no competing interest.

