## Supplementary Info for "DNAffinity: A Machine-Learning Approach to Predict DNA Binding Affinities of Transcription Factors"

---

**Supplementary Methods**

Supplementary Tables

**Supplementary Table 1** Datasets of the features and labels used in this work and corresponding post processing methods applied.

| **Labels dataset** | **Post processing methods** |
|---|---|
| uPBM (DREAM5 (41)) | Input 35-mer oligos for 66 mouse TFs.<br>Alignment and trimming using motif detected by MEME suite (k-mer, typically 12-mer). |
| gcPBM (GSE59845 and GSE44604) | gcPBM data already aligned. When detected, removal of multiple binding sites. |
| HT-SELEX (PRJEB14744) (first dataset) | Bioconductor Package SELEX<br>Removed TFs with no k-mer counts > 100 in the 0th cycle<br>Trained and tested data from the penultimate SELEX cycle |
| HT-SELEX (PRJEB29730) | Bioconductor Package SELEX |

| | |
|---|---|
| (second dataset) | Removed TFs with no k-mer counts > 100 in the 0th cycle<br>Trained and tested data from the penultimate SELEX cycle |

| **Feature dataset** | Link |
|---|---|
| Base pair parameters (average and flexibilities) and Electrostatics | MiniABC for each tetramer (1)<br>Simulations stored in BigNASim (2)<br><br>https://github.com/Jalbiti/DNAffinity |

Supplementary Table S2. Average correlation ($R^2$) and MSE for dataset (uPBM and HT-SELEX) among all the studied proteins, using cross-validation (CV) and bootstrap respectively.

| Method | Metric | CV | Bootstrap |
|---|---|---|---|
| HT-SELEX | R^2 | 0.62 ± 0.21 | 0.66 ± 0.19 |
| HT-SELEX | MSE | 0.002 ± 0.001 | 0.001 ± 0.001 |
| uPBM | R^2 | 0.63 ± 0.12 | 0.69 ± 0.17 |
| uPBM | MSE | 0.007 ± 0.009 | 0.011 ± 0.004 |

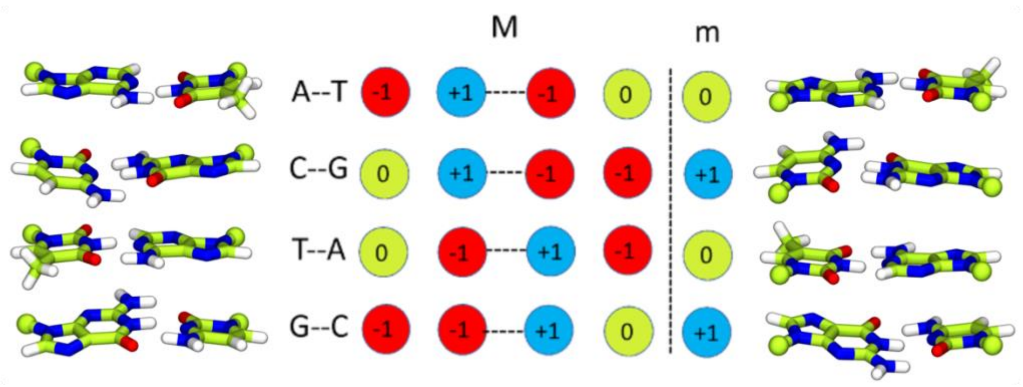


**Supplementary Figure S1.** Scheme for describing the different electrostatics of each possible base pair at the major (M) and minor (m) groove. At each position a value of +1, -1 or 0, has been assigned depending on the presence of a hydrogen bond acceptor (red), and donor (blue), or a non-polar hydrogen or methyl group (green) respectively.

**Sequence vector**

Each sequence considered as a potential TFBS was read as a k-mer vector in our algorithm. For processing, each k-mer was broken down into overlapping tetramers, and each tetramer was assigned the corresponding 4 classes of features (see Supplementary Figure S2). We considered 17 per-tetramer features and 256 presence features whose number is not dependent on the k-mer length; for example, for a 10-mer we considered a total of 375 = 17*(10-3) + 256 parameters.

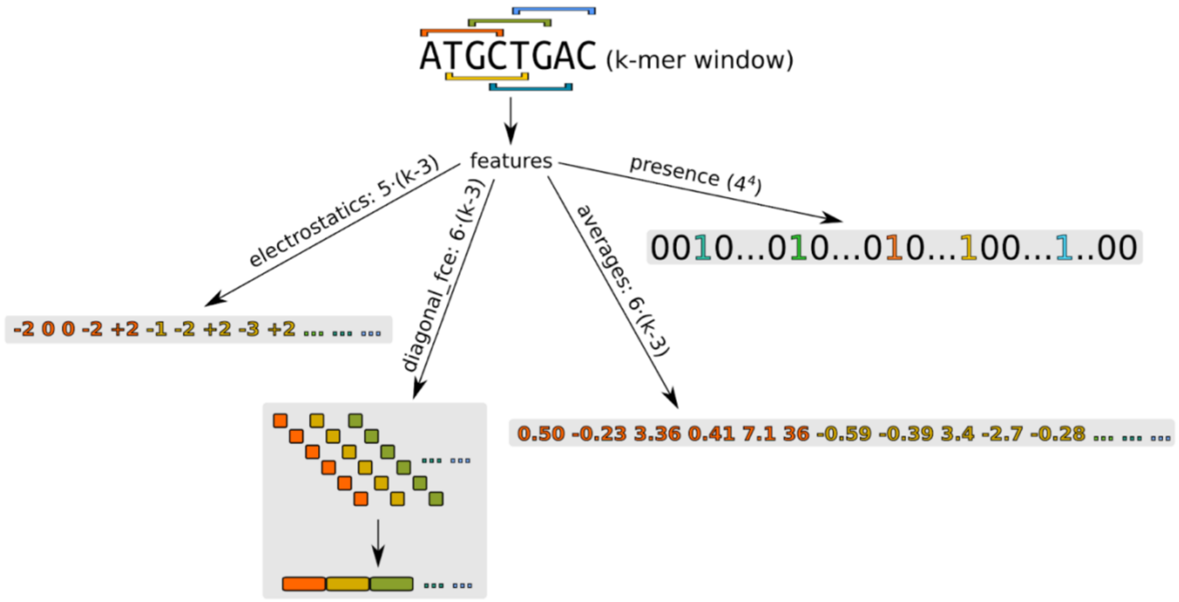


**Supplementary Figure S2.** Scheme of the sequence vector and the division into overlapping tetramers, with the attribution of each class of parameters.

**Undersampling approaches**

Undersampling was applied to the uPBM data to have a clear distinction between high and low affinity sequence without the extreme redundancy of low affinity points. More specifically, we applied undersampling only to uPBM data because in these datasets almost all TFs had only about 50-100 high-affinity k-mers (with affinities above 0.2) out of 25,000 data points, making the training extremely skewed so that the important data was lost among the big amount of low-affinity kmers (as an example, see the undersampling of the Ar dataset in Supplementary Figure S3).

To perform the undersampling, we divided the interval [0,1] of affinities into $N$ smaller subintervals, namely:

$$[0,1] = k = 0N - 1\left[\frac{k}{N}, \frac{k+1}{N}\right]$$

If $N$ is large (usually N=25000), the intervals closer to 0 will have a large density of points as compared to the intervals closer to 1. We can use this fact to pick samples from the low-affinity points and hence reduce the excess of these data points. In other words, if k ≤ N/2 we only picked $n = 1$ point from the interval $\left[\frac{k}{N}, \frac{k+1}{N}\right]$and if k > N/2 we picked all the available data in the interval, if any.

By using a revisited version of the k nearest neighbors method (https://towardsdatascience.com/machine-learning-basics-with-the-k-nearest-neighbors-algorithm-6a6e71d01761) we made sure that no range of affinities was missing and at the same time we got rid of possible noise.

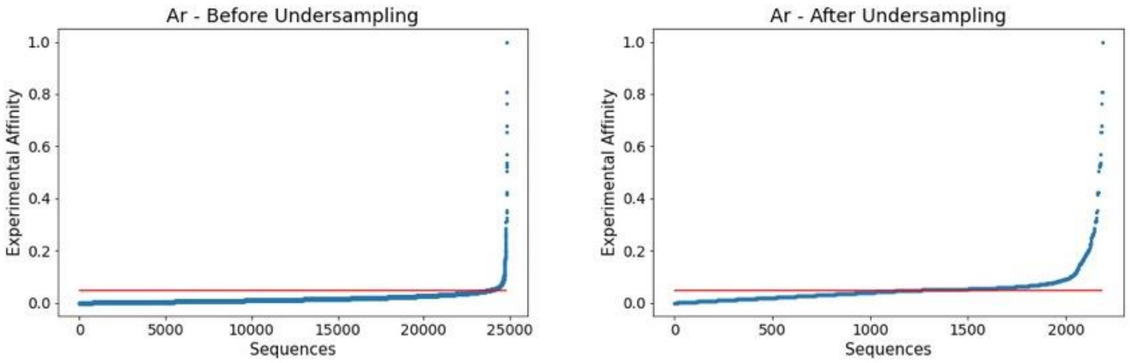


**Supplementary Figure S3**. Illustrative example of the affinity profile before (left) and after (right) undersampling of the uPBM dataset for the Ar TF. We divided the range of affinities into 25,000 intervals, and set a threshold not to undersample the top affinity 1,000 sequences, which translates into an affinity threshold around 0.05. This yielded 1196 intervals (affinity < 0.05) for which we sampled the data. In each one of these intervals we randomly take into consideration k=1 point, and from the interval above the threshold we take the remaining 1000 points. In short, we have 1,000 points to cover almost the whole range of affinity (range 0.05-1) and 1,196 of very low affinity (range 0-0.05).

For the HT-SELEX data, we removed noisy data using the 'Probability' column given by the Markov Model to under sample the dataset: we only used the top 10% points regarding probabilities to train the model and we removed the lower tail. The efficiency and reliability of this method has been previously proven (3).

**Removal of multiple binding sites**

For the gcPBM datasets (GSE59845 and GSE44604), where each probe in the array is aligned at the central TF binding site, sequences that included possible multiple binding sites were removed. For this purpose, we first generated a 6 bps PWM using the top affinity 36-mers and used it to scan the flanking sequence surrounding the central TFBS The sequence was discarded when comparing the flanking sequences with the PWM generated we found more than 4 continuous bps in common. This procedure is an adaptation of the protocol described in (4), which provides a final number of curated sequences similar to ours.

**Feature importance**

To calculate the importance of each feature, we used a function called "feature importances" integrated in the sklearn package that we used for the Random Regressor algorithm; the higher the value, the more important the feature. The importance of a feature is computed as the (normalized) total reduction of the criterion brought by that feature. It is also known as the Gini importance (https://towardsdatascience.com/the-mathematics-of-decision-trees-random-forest-and-feature-importance-in-scikit-learn-and-spark-f2861df67e3).

**HT-SELEX data quality assessment**

Analysis is based on HT-SELEX raw observed counts. We were able to classify whether the raw data would allow us to predict the binding affinities with high accuracy, using the correlation between the counts across the different cycles and the statistical distribution across the penultimate cycle, which was used for prediction. The SVM was then trained to classify all proteins, analysing whether our DNA protein-affinity predictor would perform better than a random predictor ($R^2 > 0.5$) and all proteins were reassessed using this classification as a filter. A radial-basis function kernel (5), a regularization parameter of 10 and a gamma value of 0.1 were used to define our SVM parameters.

**Prediction quality**

For the $R^2$, we used the r2_score function in scikit-learn that computes the coefficient of determination; the formula used is the following:

$$R^2(y, \hat{y}) = 1 - \frac{\sum_{i=1}^{n}(y_i - \hat{y}_i)^2}{\sum_{i=1}^{n}(y_i - \bar{y})^2}$$

Where $\hat{y}_i$ is the predicted value of the i-th sample, $y_i$ is the corresponding true value for total n samples and $\bar{y}$ is the average value.

For MSE we used the mean_squared_error function in scikit-learn that computes the coefficient of determination; the formula used is the following:

$$MSE(y,\hat{y}) = \frac{1}{n_{samples}} \sum_{i=0}^{n_{samples}-1} (y_i - \hat{y}_i)^2$$

Where $\hat{y}_i$ is the predicted value of the i-th sample, $y_i$ is the corresponding true value for total n samples.

For Pearson correlation we used the stats.pearsonr function in scikit-learn (https://docs.scipy.org/doc/scipy/reference/generated/scipy.stats.pearsonr.html), the formula used is the following:

$$r = \frac{\sum_{i=1}^{n}(x_i - \overline{x}) \quad (y_i - \overline{y})}{\sqrt{\sum_{i=1}^{n}(x_i - \overline{x})^2 \sum_{i=1}^{n}(y_i - \overline{y})^2}}$$

Where $y_i$ is the predicted value of the i-th sample, $x_i$ is the corresponding true value for total n samples, $\overline{x}$ and $\overline{y}$are the respective mean values of the true and predicted samples.

We also tested the choice of the training set using cross-validation, which performs the simulation K times dividing the data into K partitions and using each time one different partition as a test set. Changing the algorithm to k-fold cross-validation **(K=10; 90/10 random split**), we found very similar results are obtained (see Supplementary Table S2).

### Supplementary Results

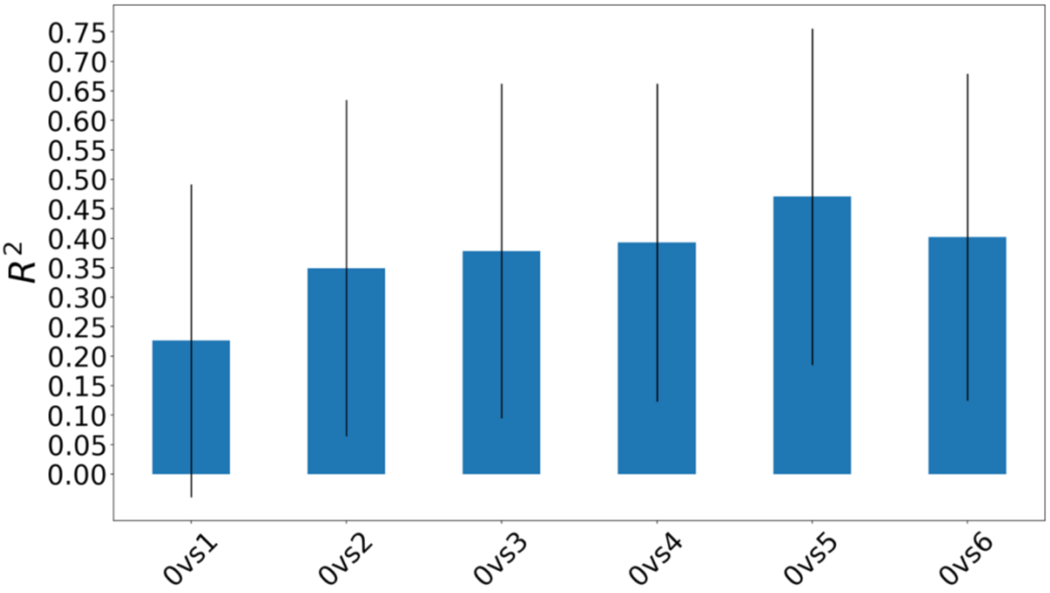


**Supplementary Figure S4.** Average correlation between experimental and predicted affinities when training on different cycle pairs among all the studied proteins, with standard deviations for the second HT-SELEX dataset (see Methods and Supplementary Table S1).

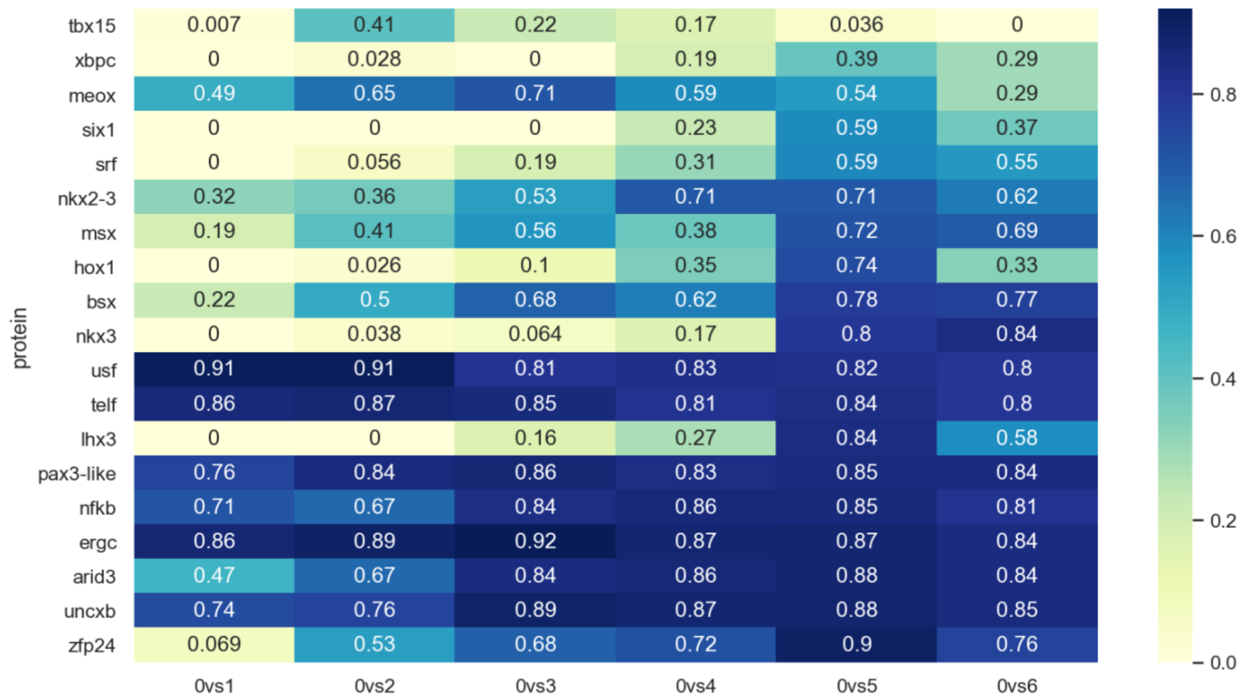


**Supplementary Figure S5.** Correlations between experimental affinities and ML predictions when model was trained on different pairs of cycles (0vs) from HT-SELEX for the second HT-SELEX dataset (see Methods and Supplementary Table S1).

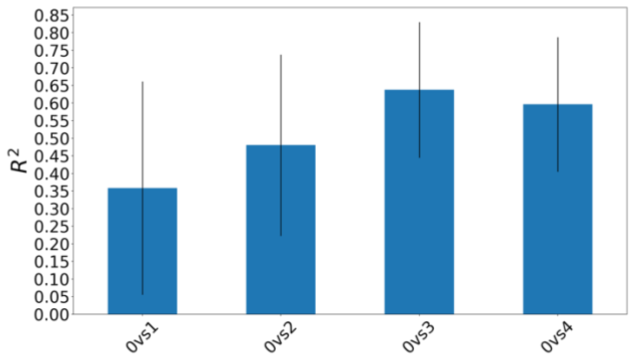

**Supplementary Figure S6.** Figure. Average correlation between experimental and predicted affinities when training on different cycle pairs among all the studied proteins, with standard deviations for the first HT-SELEX dataset (see Methods and Supplementary Table S1).

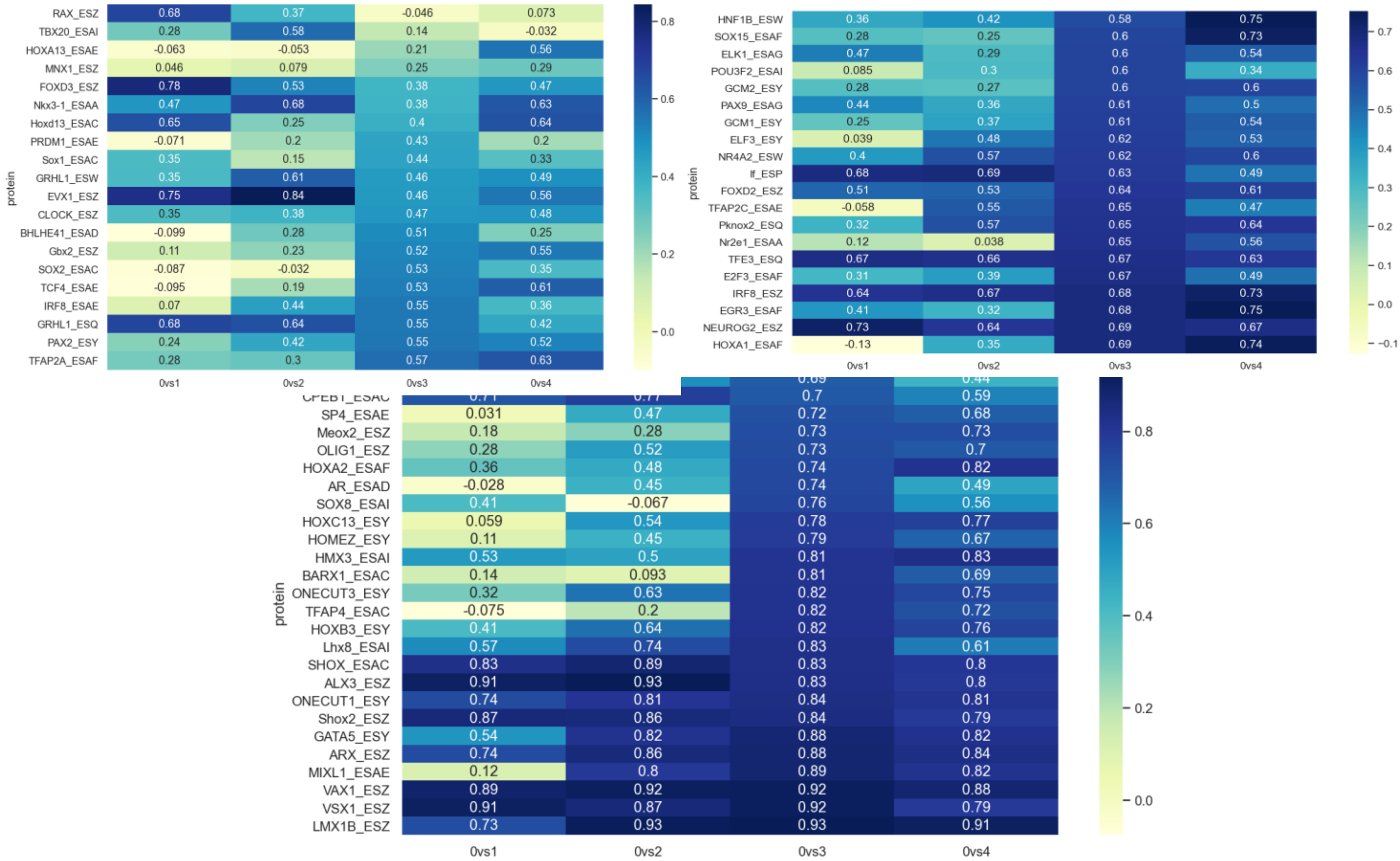


**Supplementary Figure S7.** Correlations between experimental affinities and ML predictions when model was trained on different pairs of cycles (0vs) from HT-SELEX and predicted affinity for the first dataset (see Methods and Supplementary Table S1).

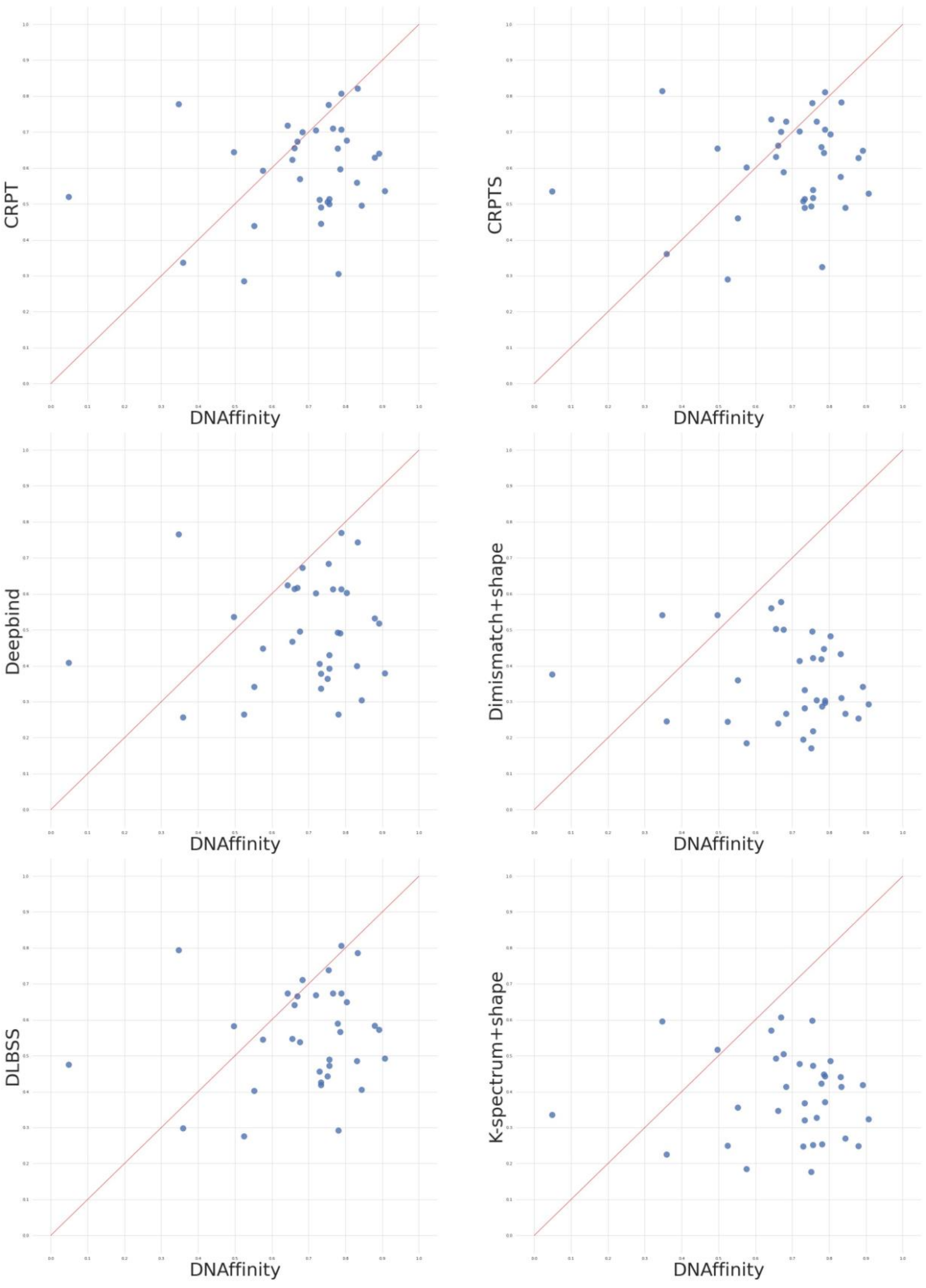


**Supplementary Figure S8.** Correlation between determination coefficients obtained by previously developed predictors and our method (DNA affinity). Methods used for benchmarking are: CRPTS/CRPT (a hybrid convolutional recurrent neural network (CNN/RNN) combining DNA sequence and DNA shape features)(6); Deepbind (CNN model based on primary DNA sequences) (7); two kernel-based methods (spectrum + shape kernel, di-mismatch + shape kernel) (8); and a deep learning method DLBSS (9). The data were taken from (6).

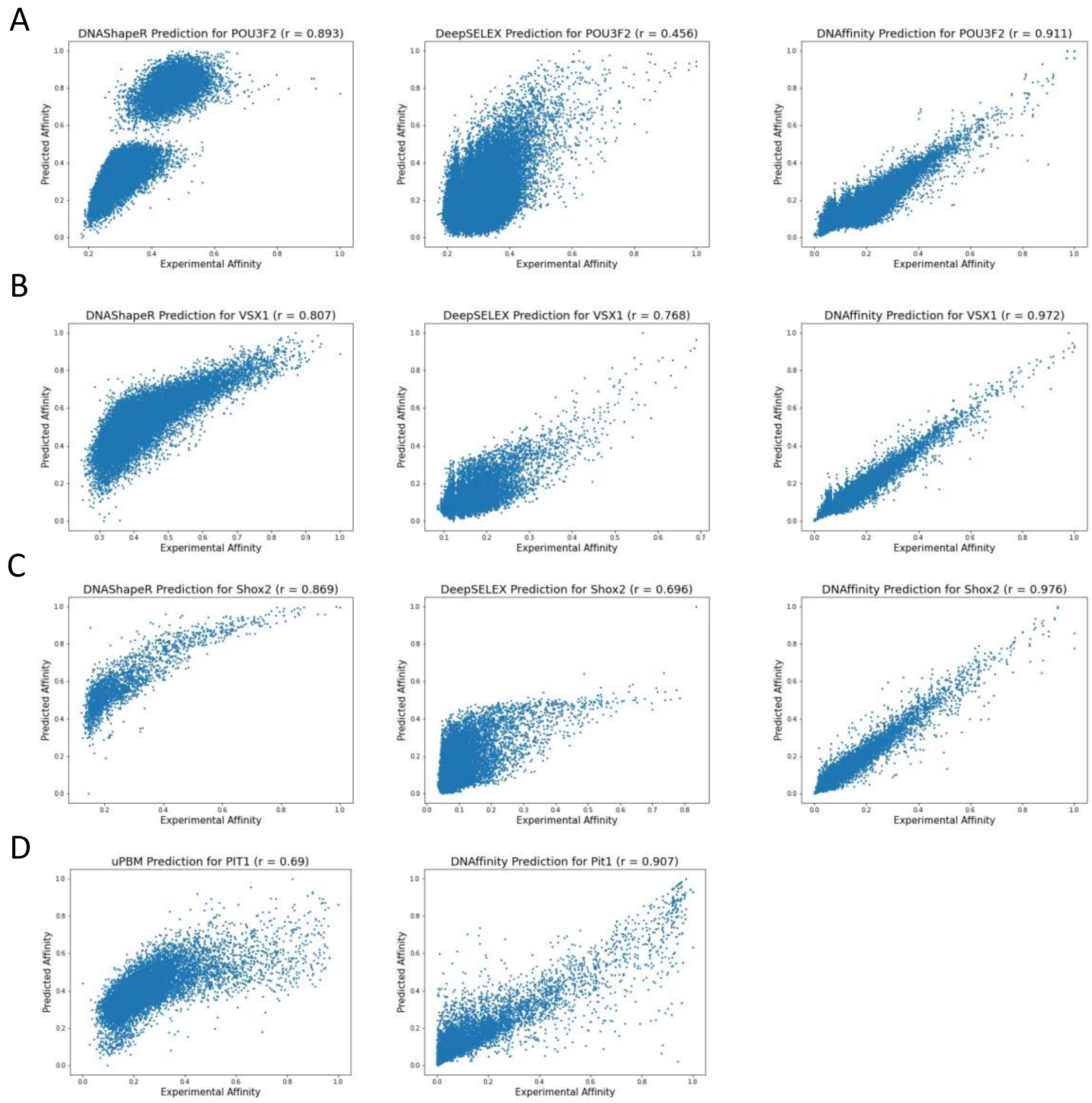


**Supplementary Figure S9. Comparison of the affinity values predicted for 3 TFs** using: A-C) DNAShapeR (left panels, (10)), DEEPSELEX (central panels, (11)) and our predictor (DNAffinity, right panel). The 3 cases selected are among all the ones studied using HT-SELEX data, for which the predictions of DNAShapeR (A), DEEPSELEX (B), DNAffinity (C) perform at its best respectively. D) CRPTS/CRPT (left panel) and DNAffinity (right panel) for a TF among all the ones studied using uPBM data with high performance.

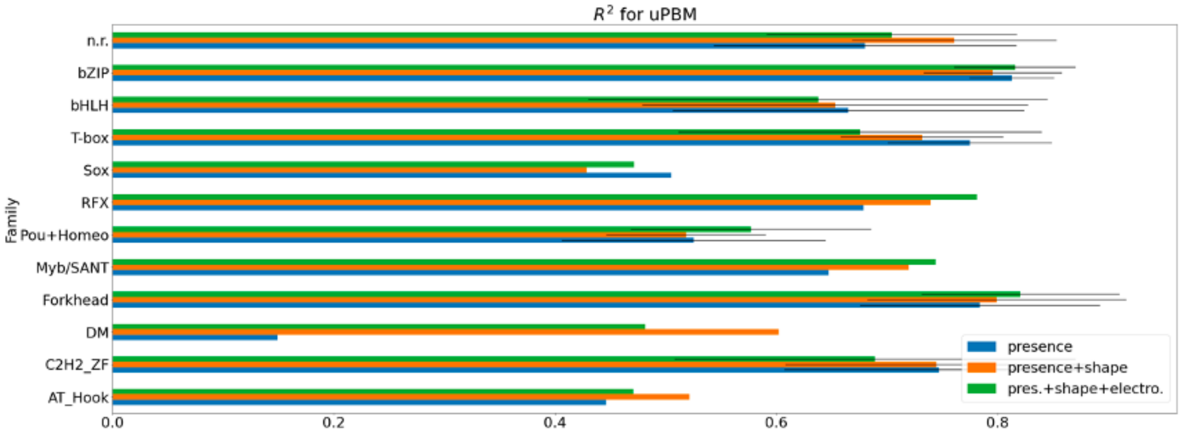


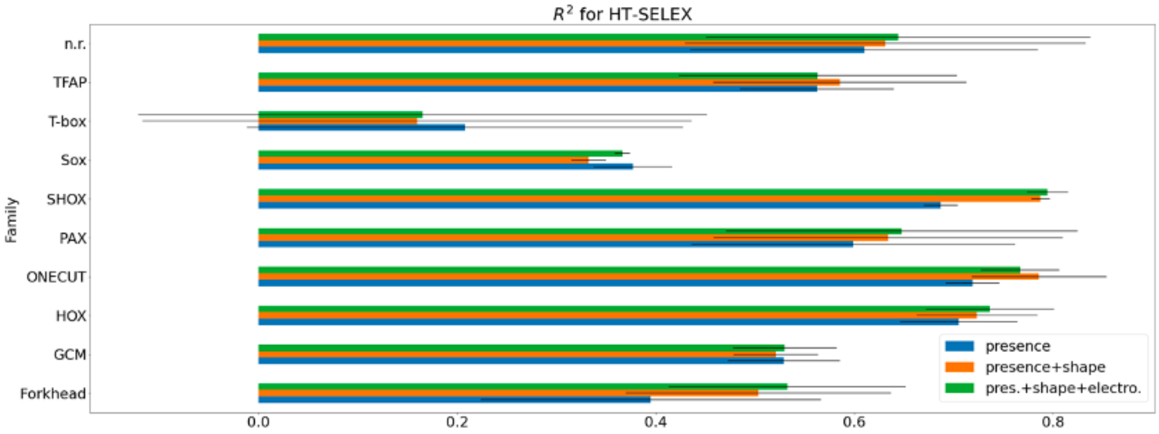


**Supplementary Figure S10.** Average correlations by protein families, with standard deviation, using three different combinations of features for uPBM: presence (in blue), base pair parameters (presence + shape, in orange) and all combined (presence+shape+electrostatics, in green).

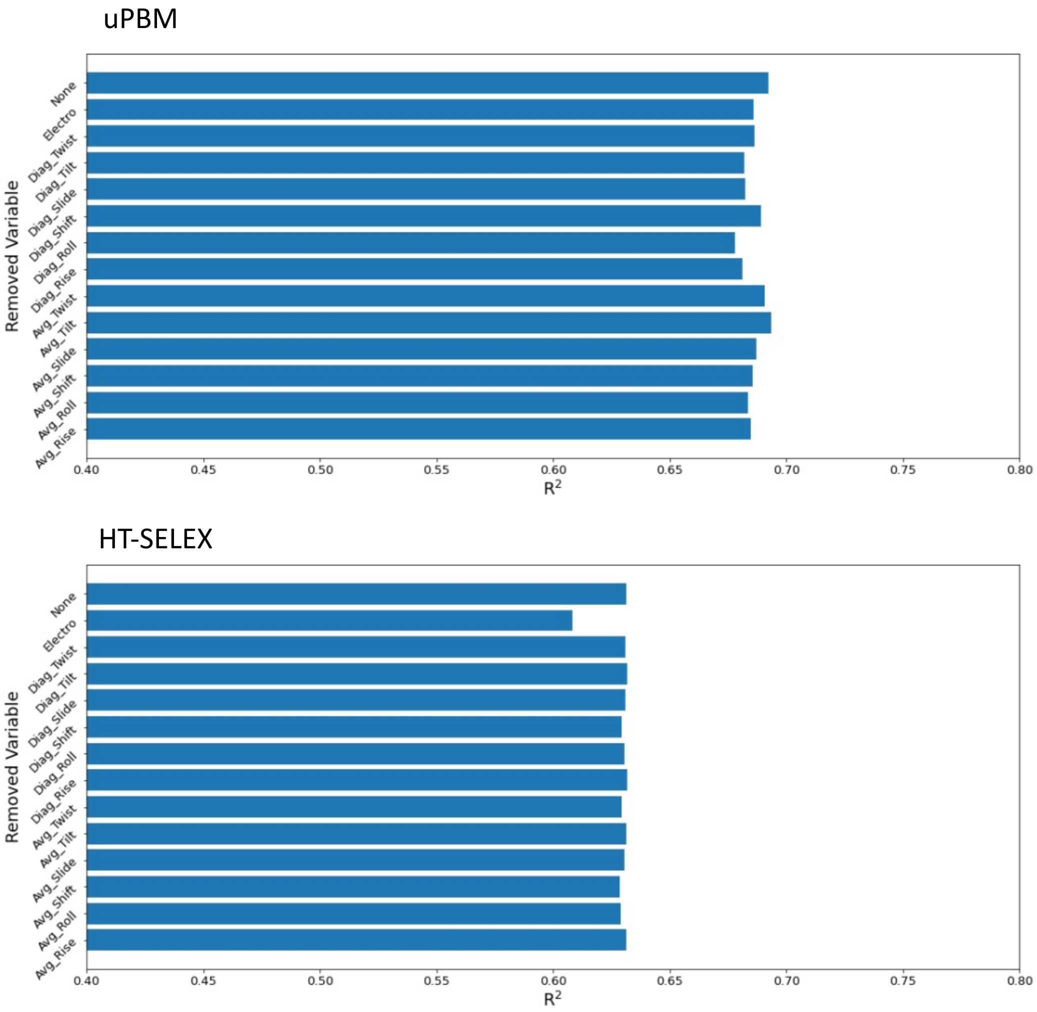


**Supplementary Figure S11.** Determination coefficient ($R^2$) between predicted and experimental affinities using uPBM (top panel) and first HT-SELEX (bottom panel) dataset, considering all the descriptors used in the machine learning algorithm (none) and removing each variable one by one (average, stiffness parameters and electrostatics).

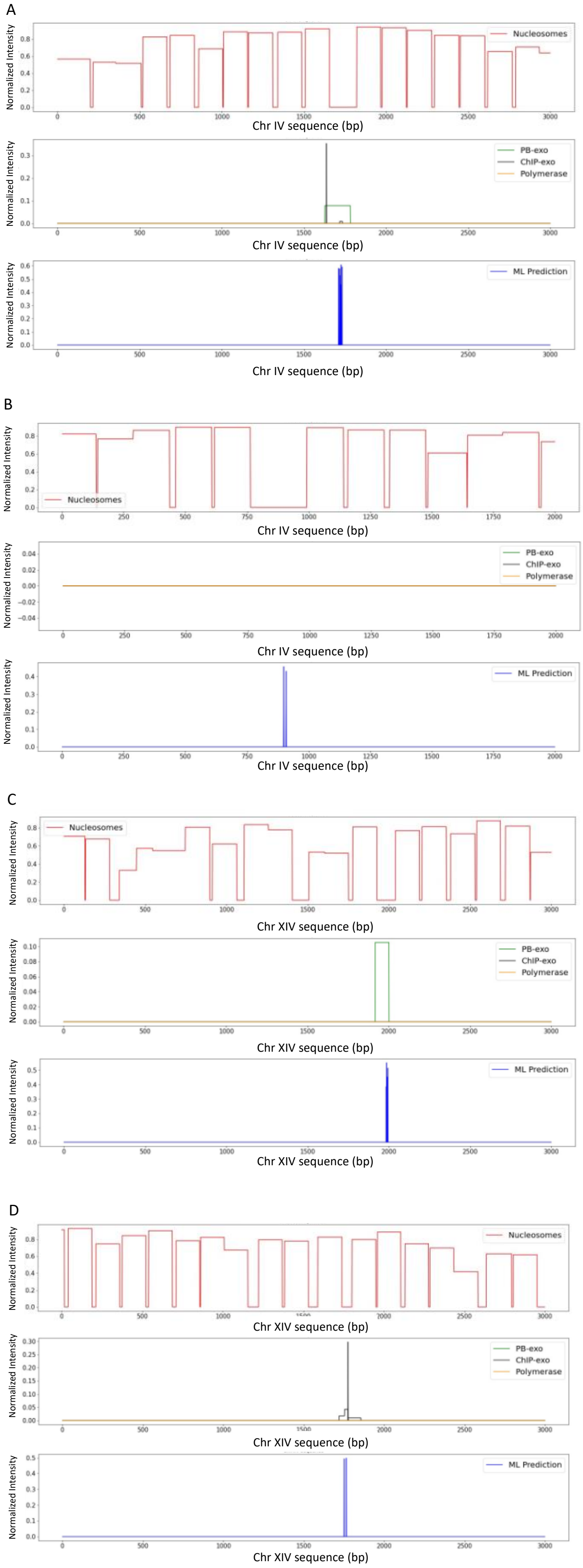
A
B
C
D
Normalized Intensity
Nucleosomes
PB-exo
ChIP-exo
Polymerase
ML Prediction
Chr IV sequence (bp)
Chr XIV sequence (bp)

**Supplementary Figure S12.** Examples of the TF positioning prediction compared with experimental data (CHiP-exo, PB-exo and Polymerase; green, black and yellow lines, respectively) and nucleosome positioning (red line). A) True positive case, the ML prediction coincides with experimentally detected TFBS in a nucleosome free region. B) False positive case, the ML prediction coincides with a nucleosome free region, but no TFBS has been detected experimentally. C-D) ”Contentious” false positive case, the ML prediction coincides with a nucleosome free region and a TFBS only detected by one experimental technique (PB-exo and Chip-exo respectively).

### Supplementary Tables

**Supplementary Table S3.** LOGOs of the position weight matrices after the training for uPBM and HT-SELEX for each TF.

| **LOGO - uPBM dataset** |
|---|
| 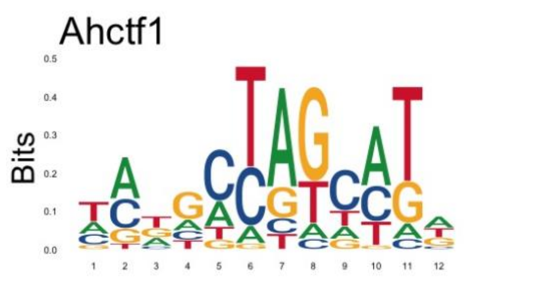  |
| 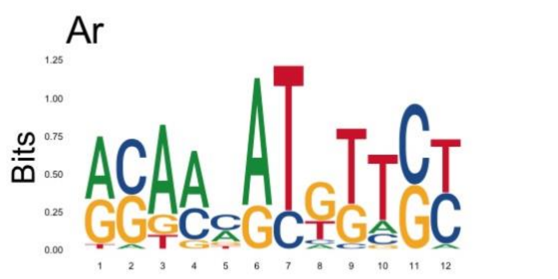  |
| 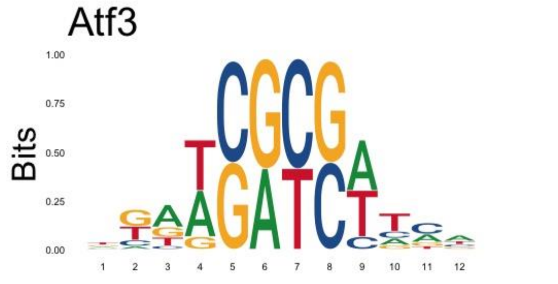  |
| 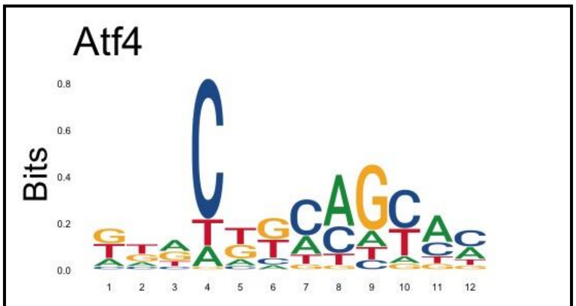  |
| 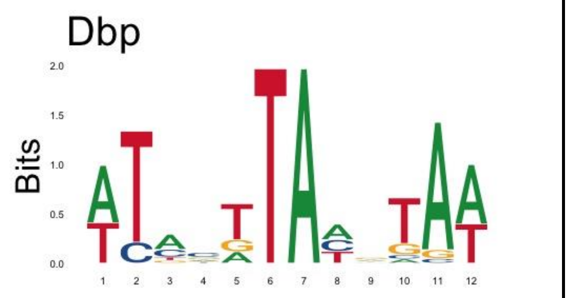  |
| 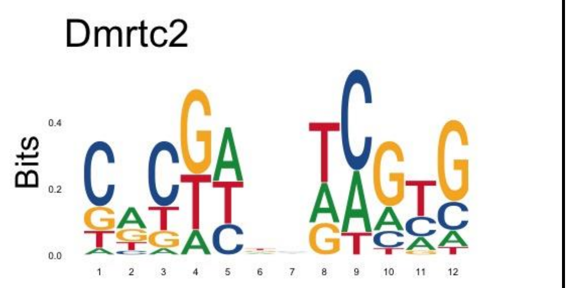  |

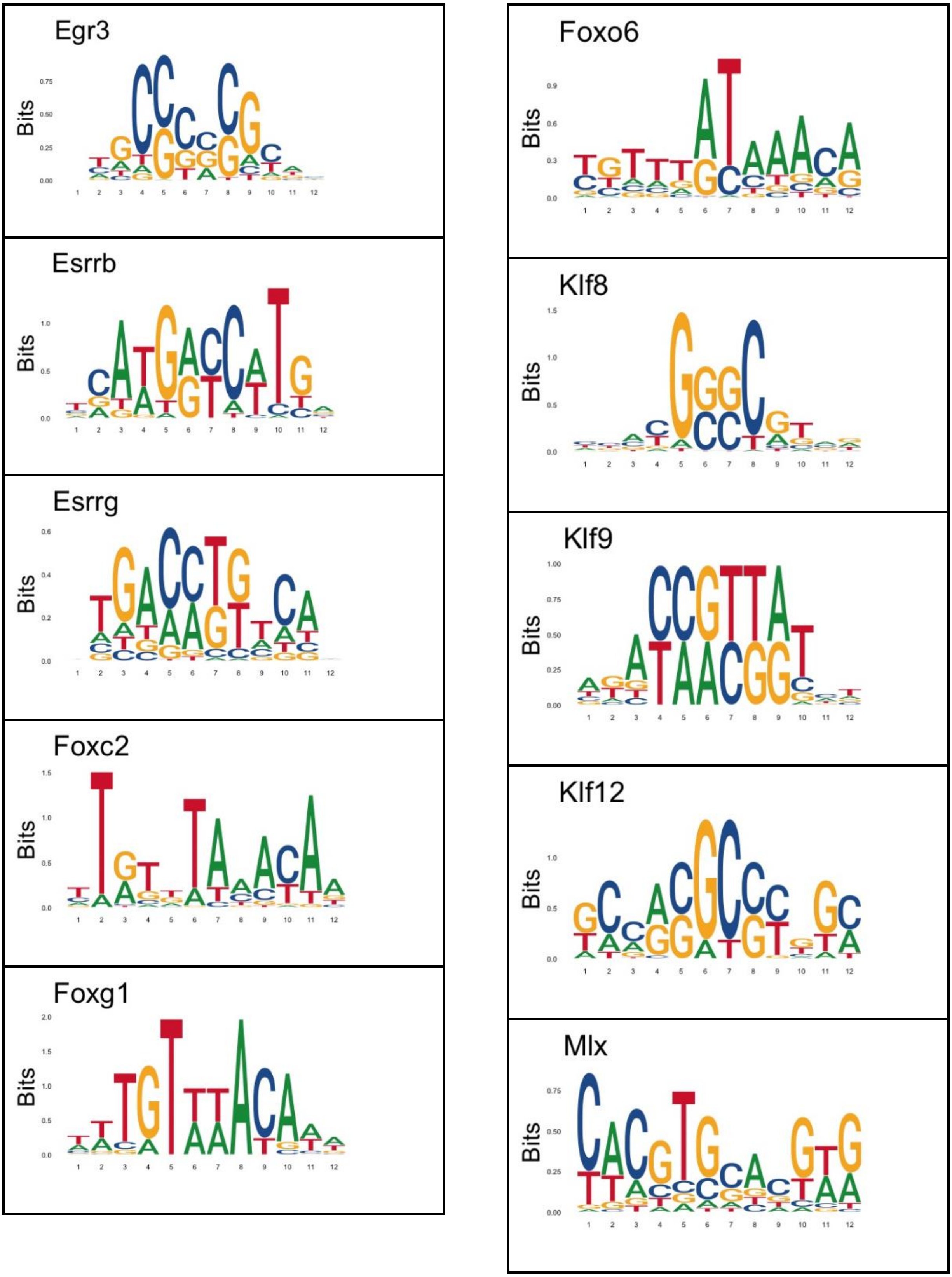

Egr3
Bits
Esrrb
Bits
Esrrg
Bits
Foxc2
Bits
Foxg1
Bits
Foxo6
Bits
Klf8
Bits
Klf9
Bits
Klf12
Bits
Mlx
Bits

Mybl2

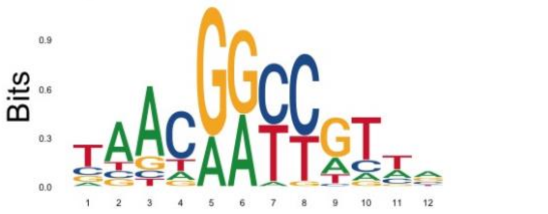

Nr2f1

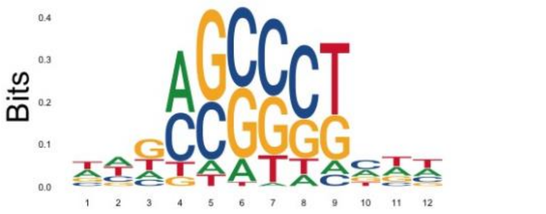

Mzf1

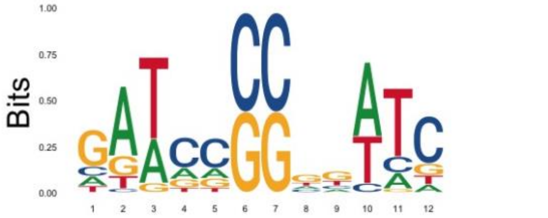

Nr2f6

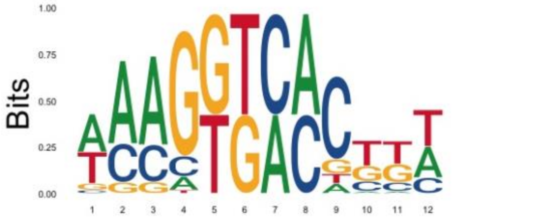

Nfil3

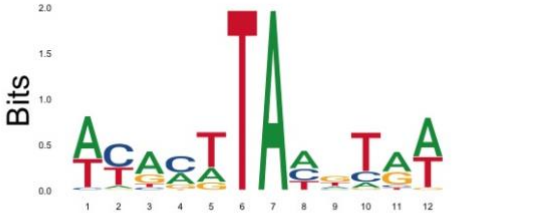

Nr4a2

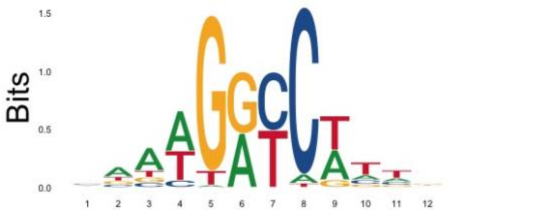

Nhlh2

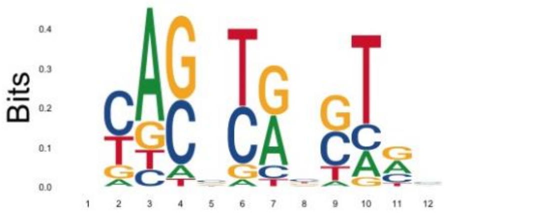

Nr5a2

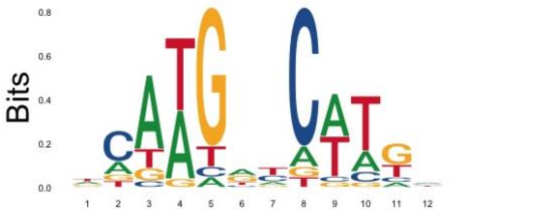

Nkx2-9

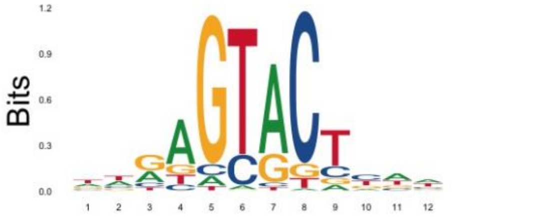

P42pop

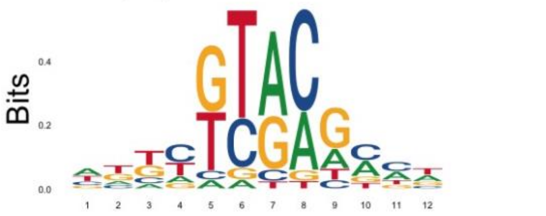

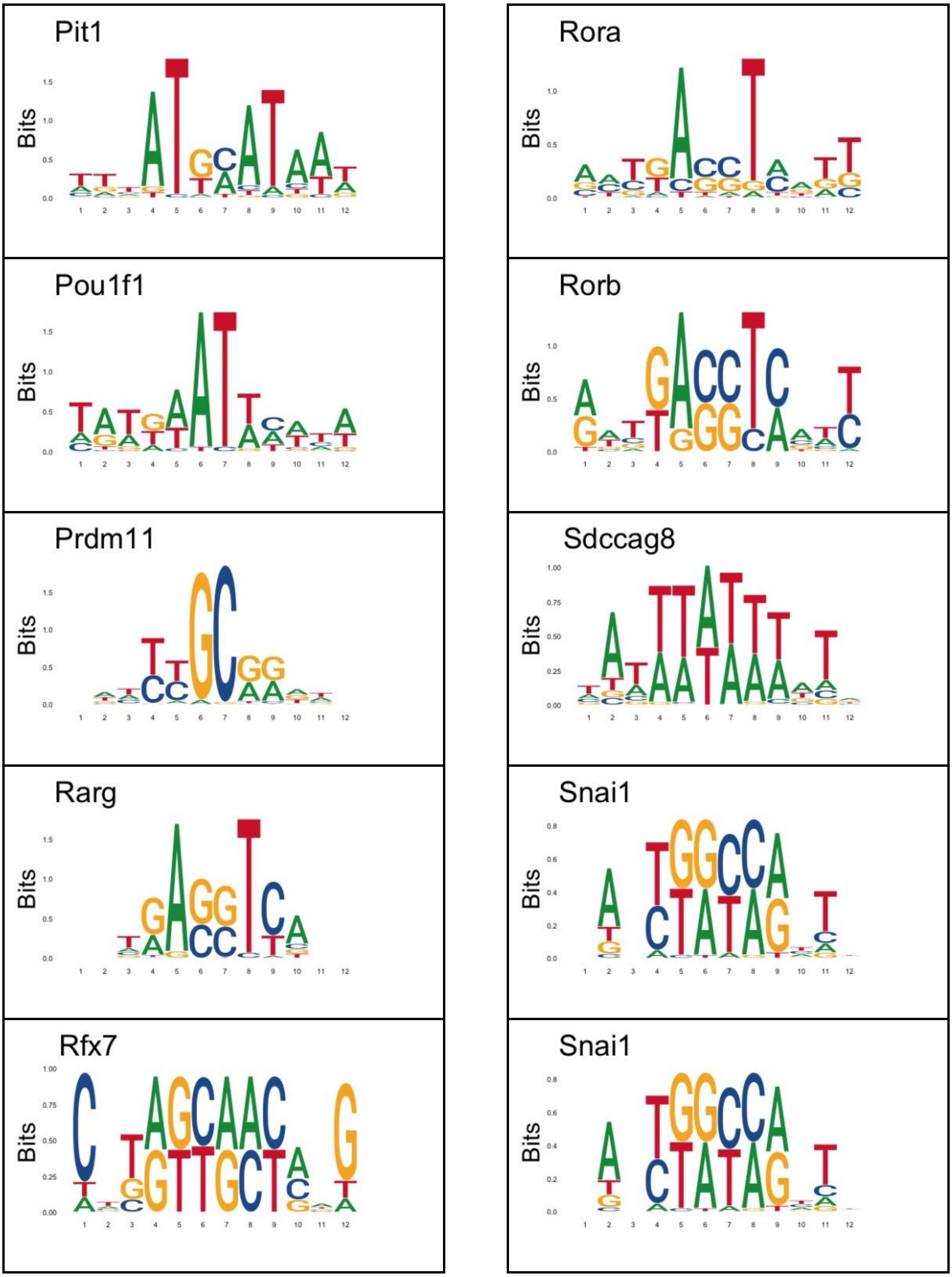
Pit1
Rora
Pou1f1
Rorb
Prdm11
Sdccag8
Rarg
Snai1
Rfx7
Snai1
Bits

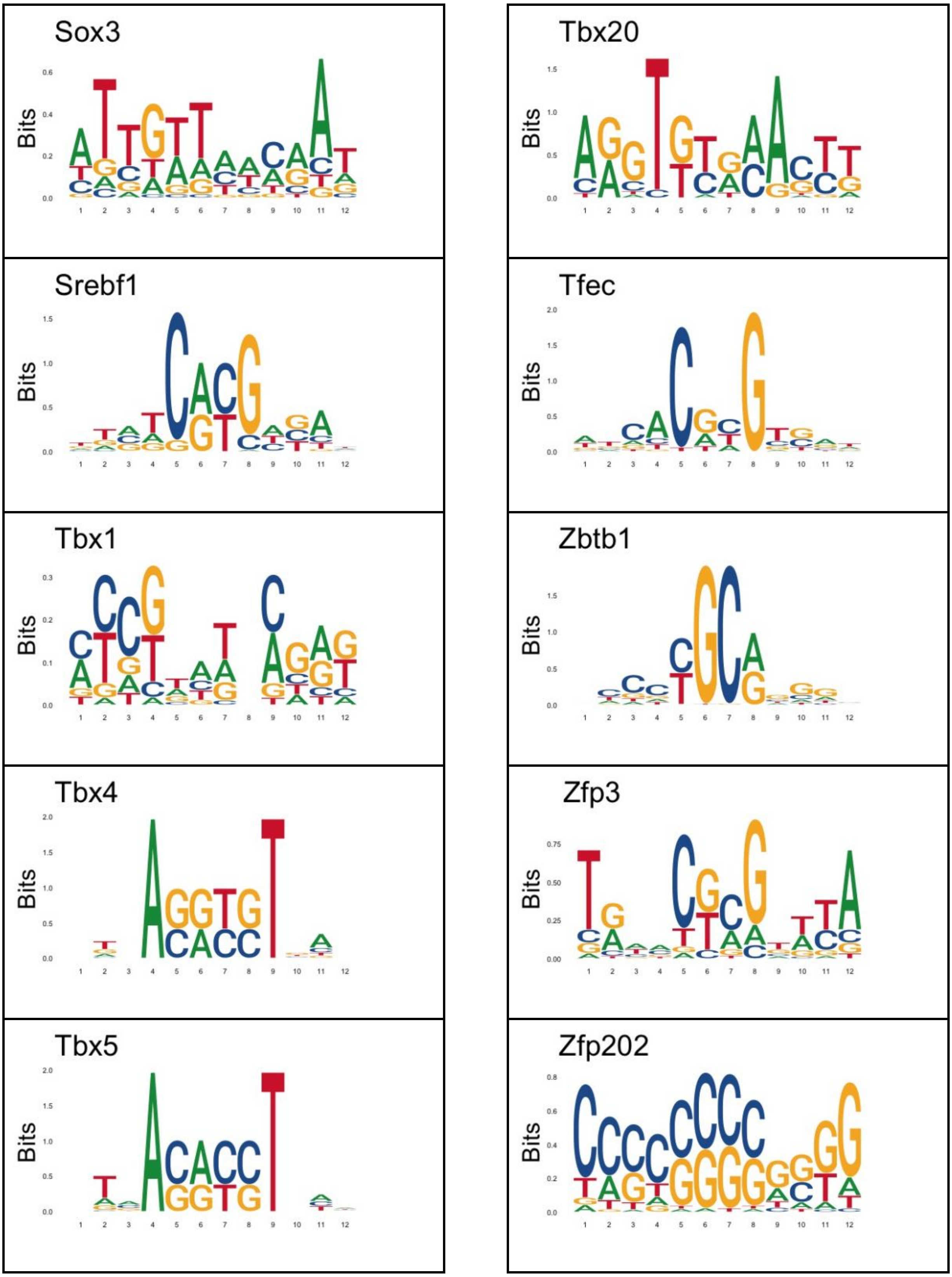
Sox3
Tbx20
Srebf1
Tfec
Tbx1
Zbtb1
Tbx4
Zfp3
Tbx5
Zfp202
Bits

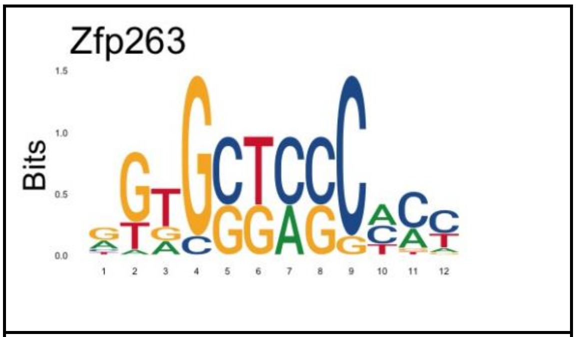
Zfp263
Bits

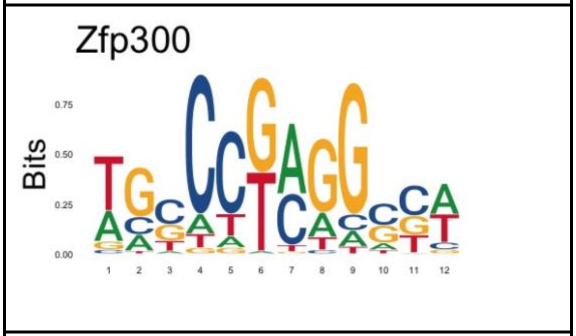
Zfp300
Bits

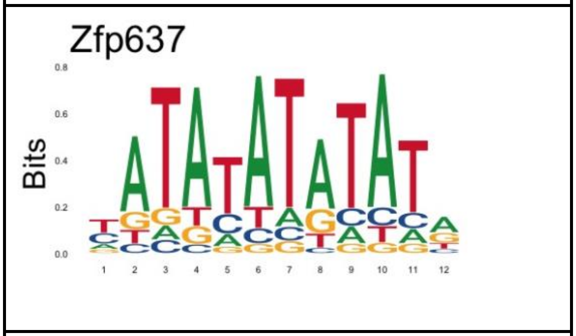
Zfp637
Bits

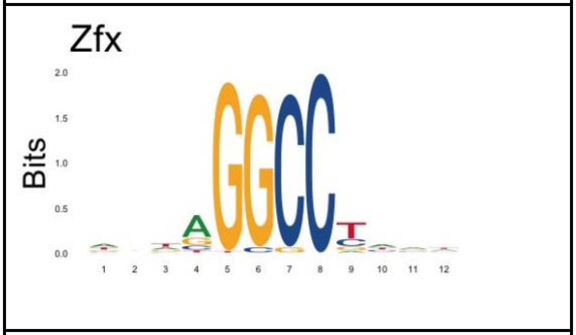
Zfx
Bits

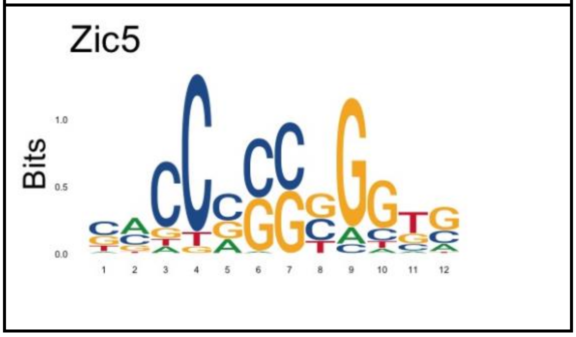
Zic5
Bits

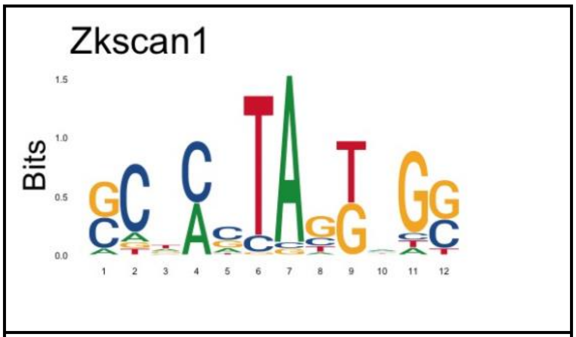
Zkscan1
Bits

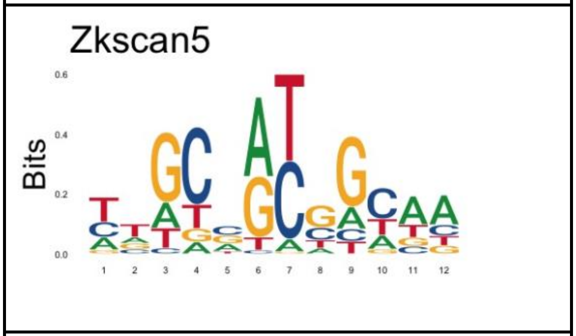
Zkscan5
Bits

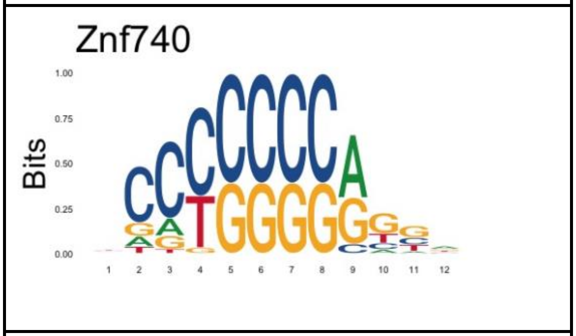
Znf740
Bits

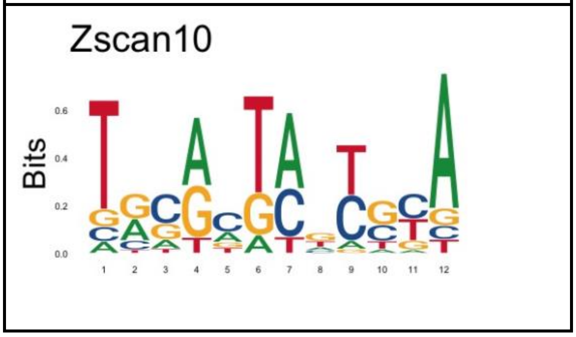
Zscan10
Bits

**LOGO – HT-SELEX dataset**

ALX3

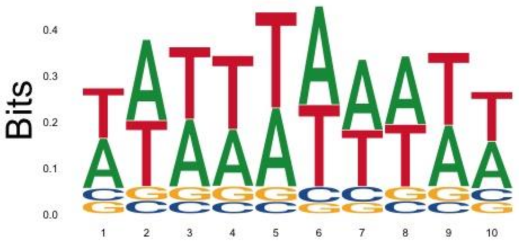

AR

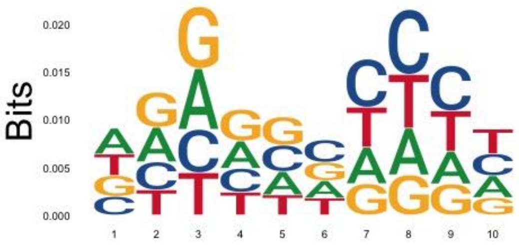

arid3

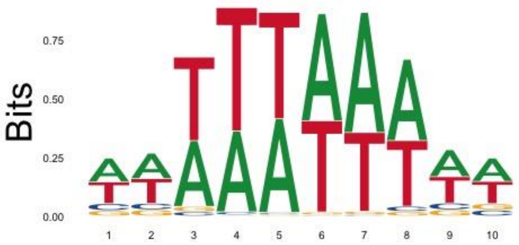

ARX

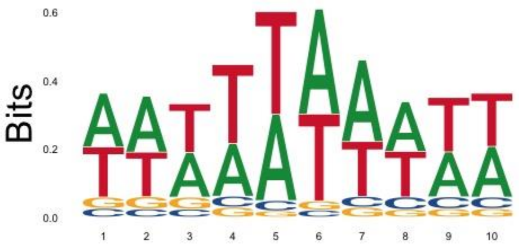

BARX1

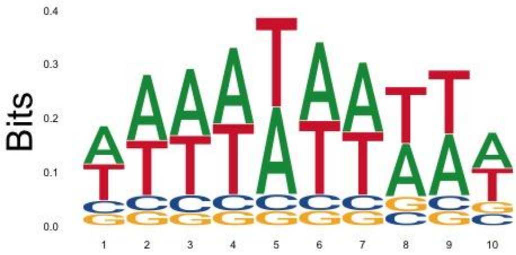

BHLHE41

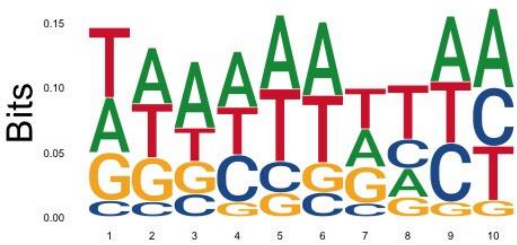

bsx

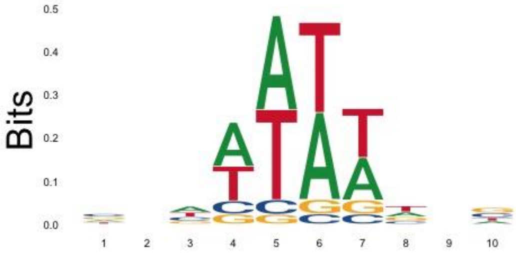

CLOCK

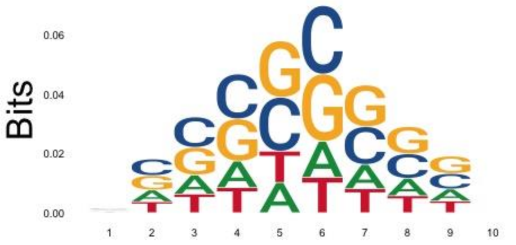

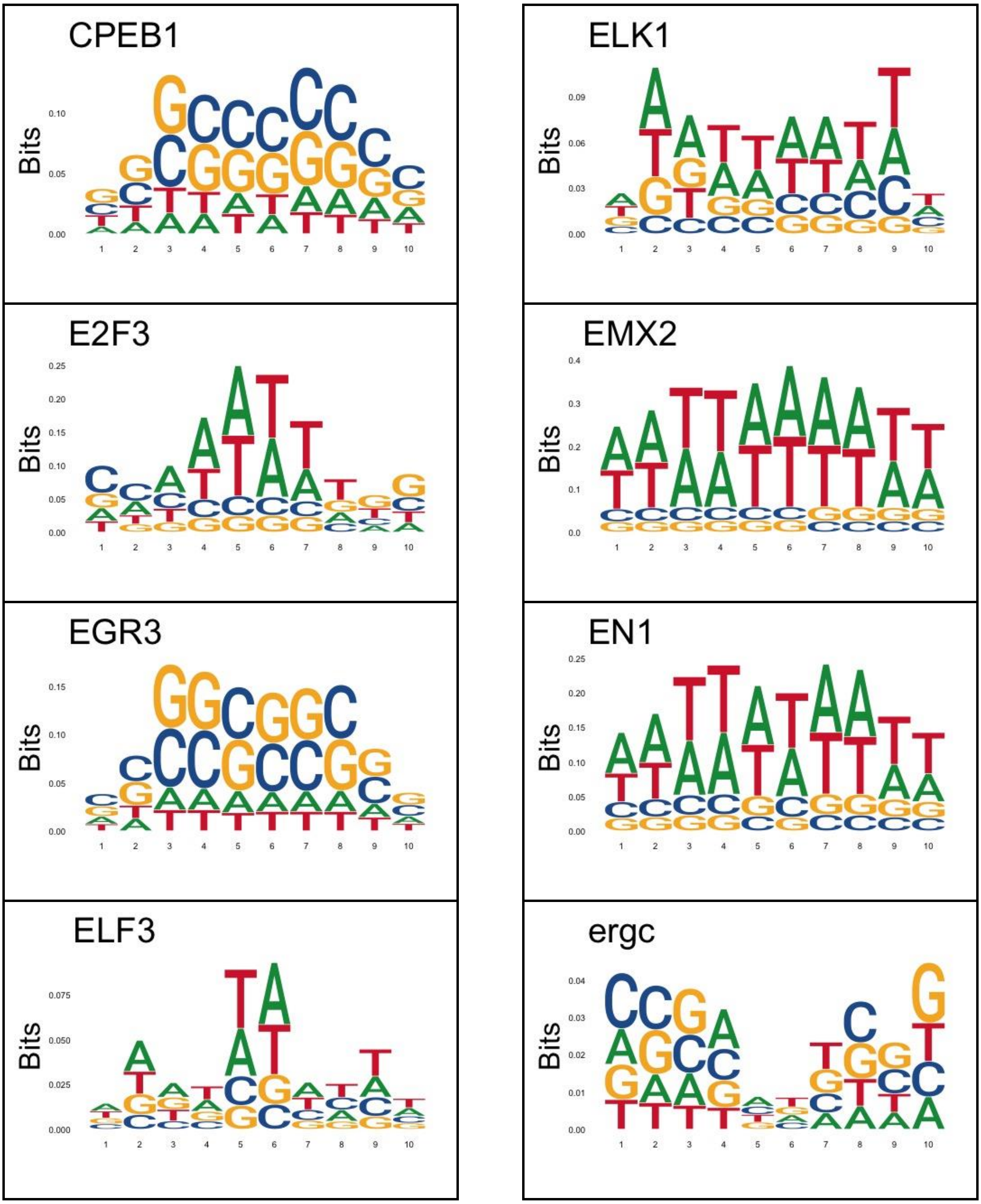
CPEB1
ELK1
E2F3
EMX2
EGR3
EN1
ELF3
ergc
Bits

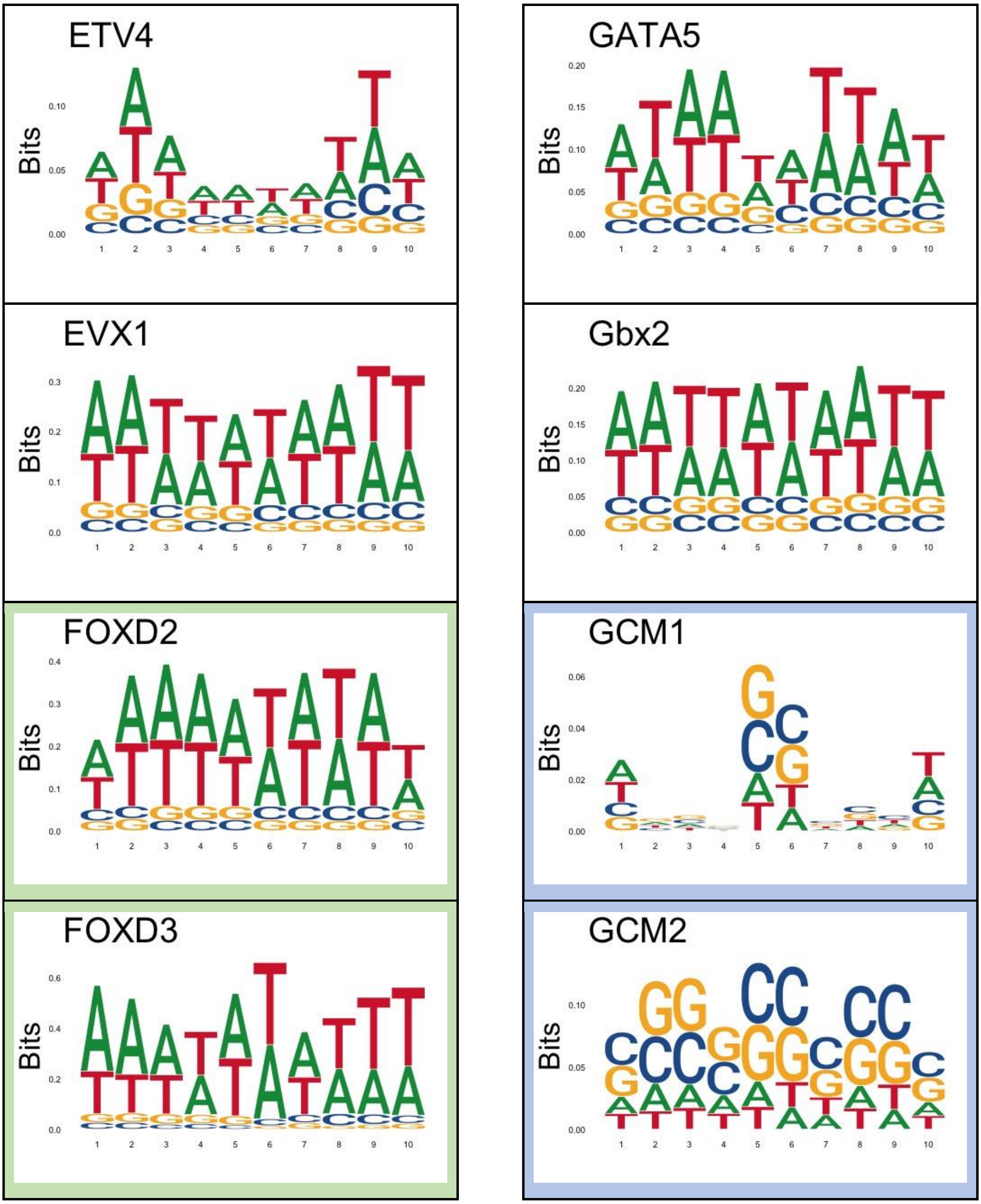
ETV4
GATA5
EVX1
Gbx2
FOXD2
GCM1
FOXD3
GCM2
Bits

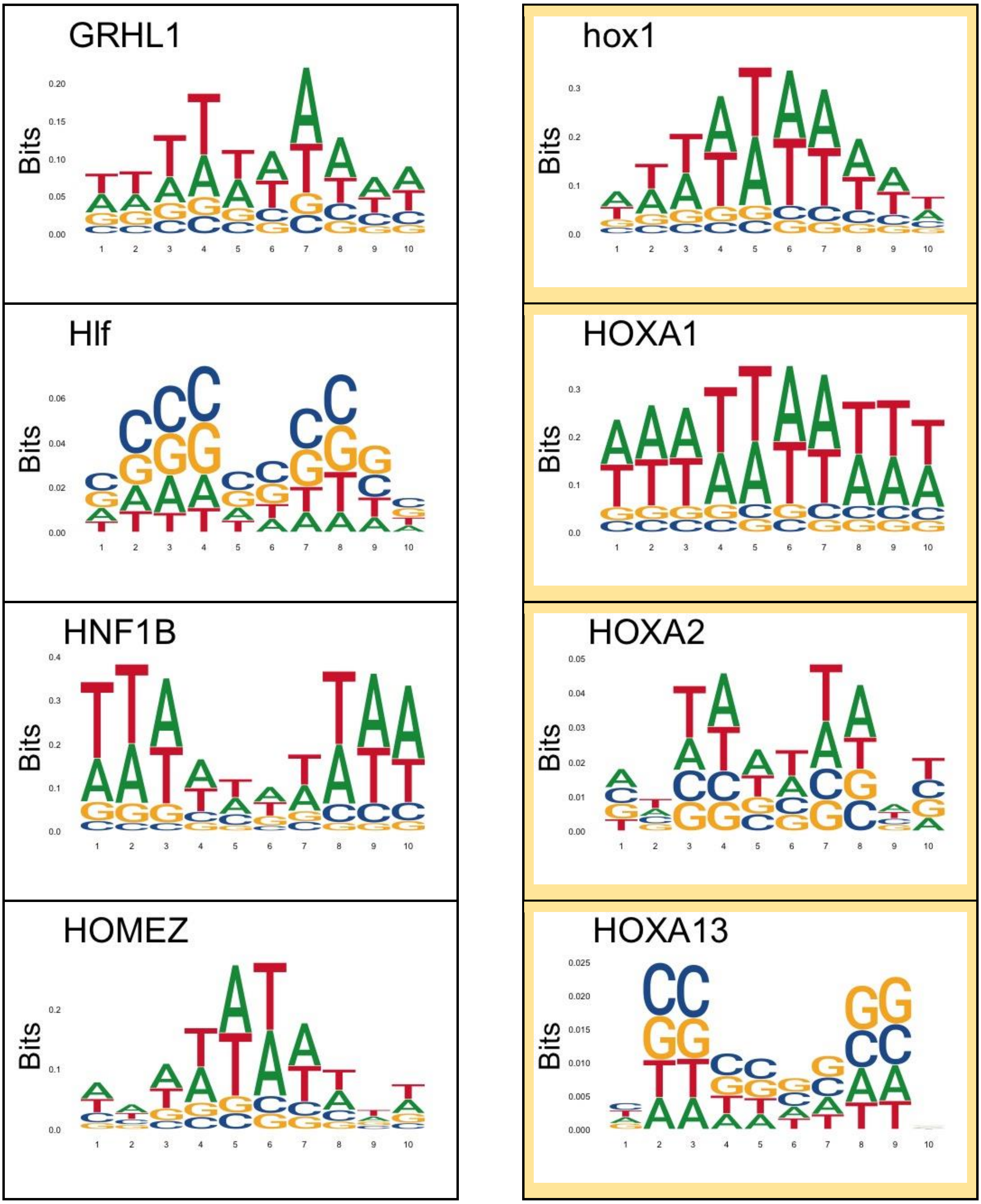
GRHL1
hox1
Hlf
HOXA1
HNF1B
HOXA2
HOMEZ
HOXA13
Bits

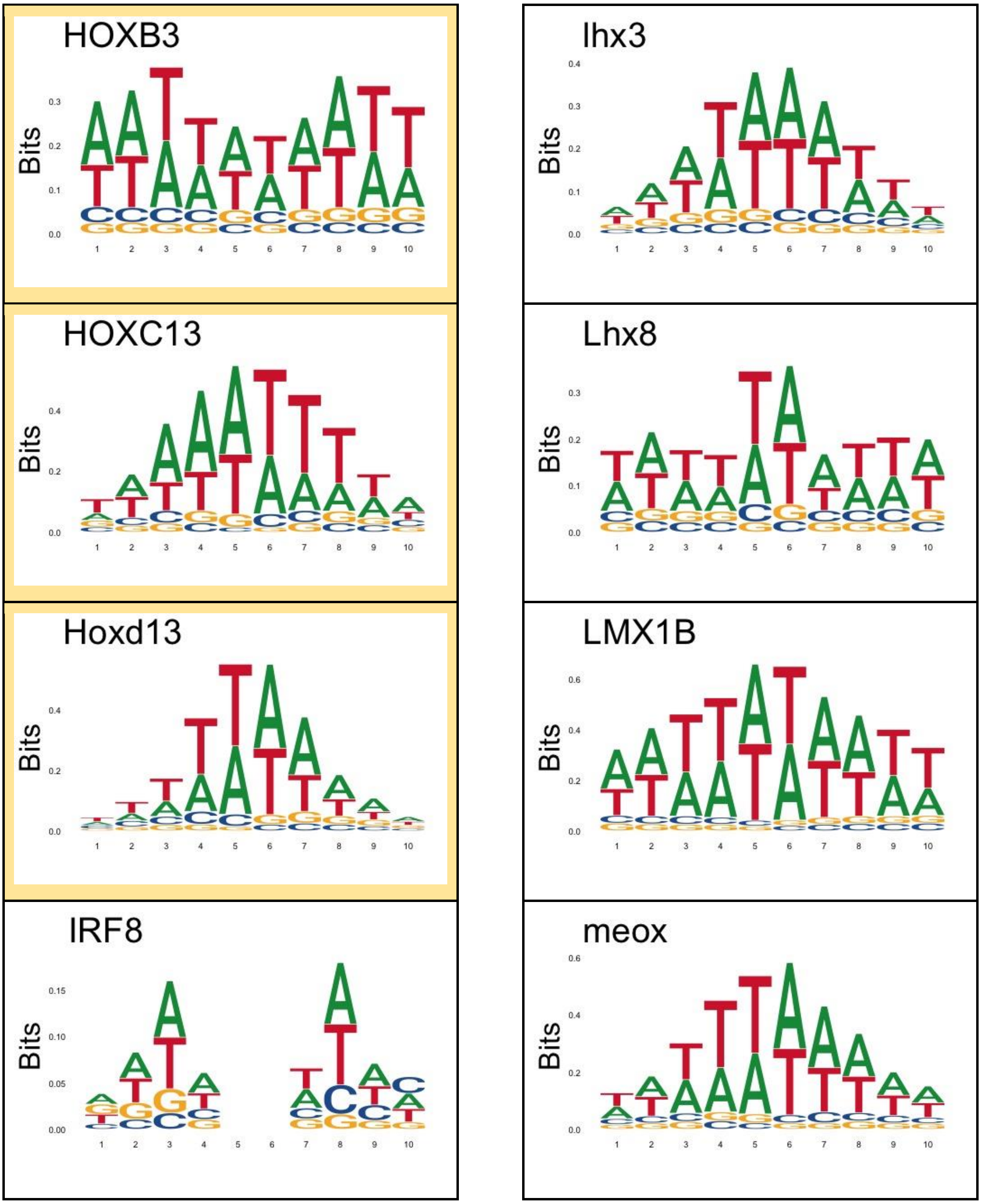
HOXB3
lhx3
HOXC13
Lhx8
Hoxd13
LMX1B
IRF8
meox
Bits
1 2 3 4 5 6 7 8 9 10

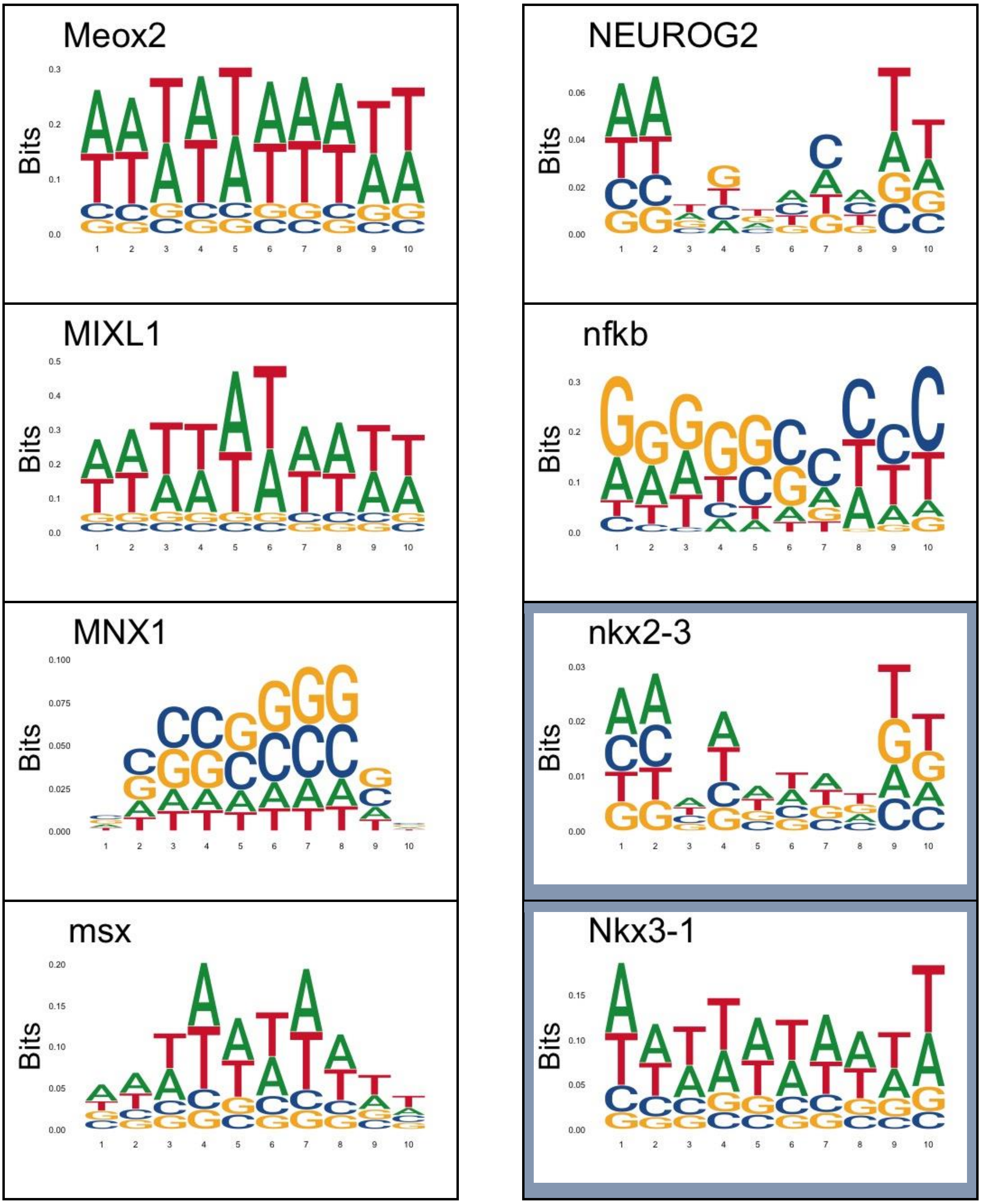
Meox2
NEUROG2
MIXL1
nfkb
MNX1
nkx2-3
msx
Nkx3-1
Bits

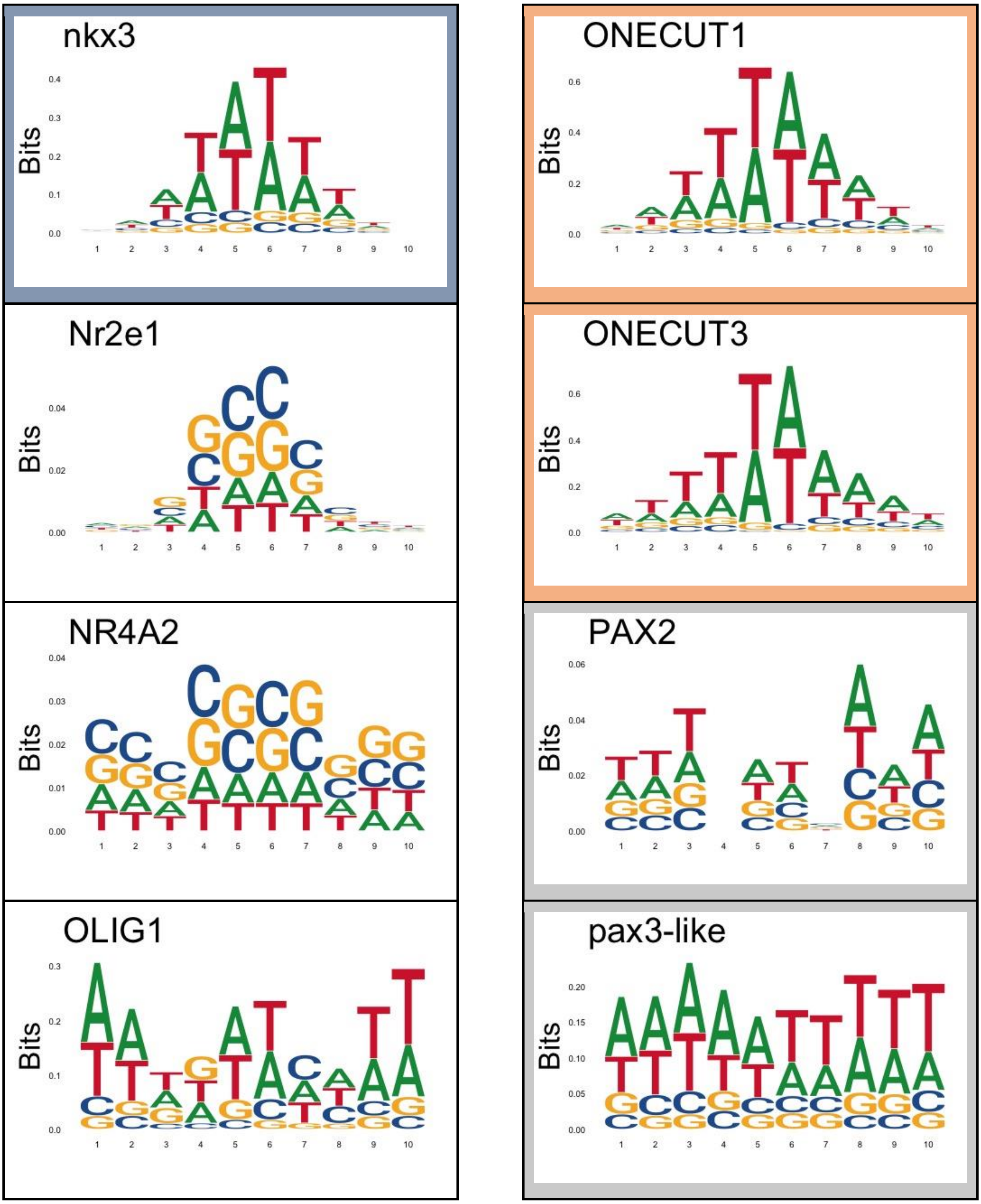
nkx3
ONECUT1
Nr2e1
ONECUT3
NR4A2
PAX2
OLIG1
pax3-like
Bits

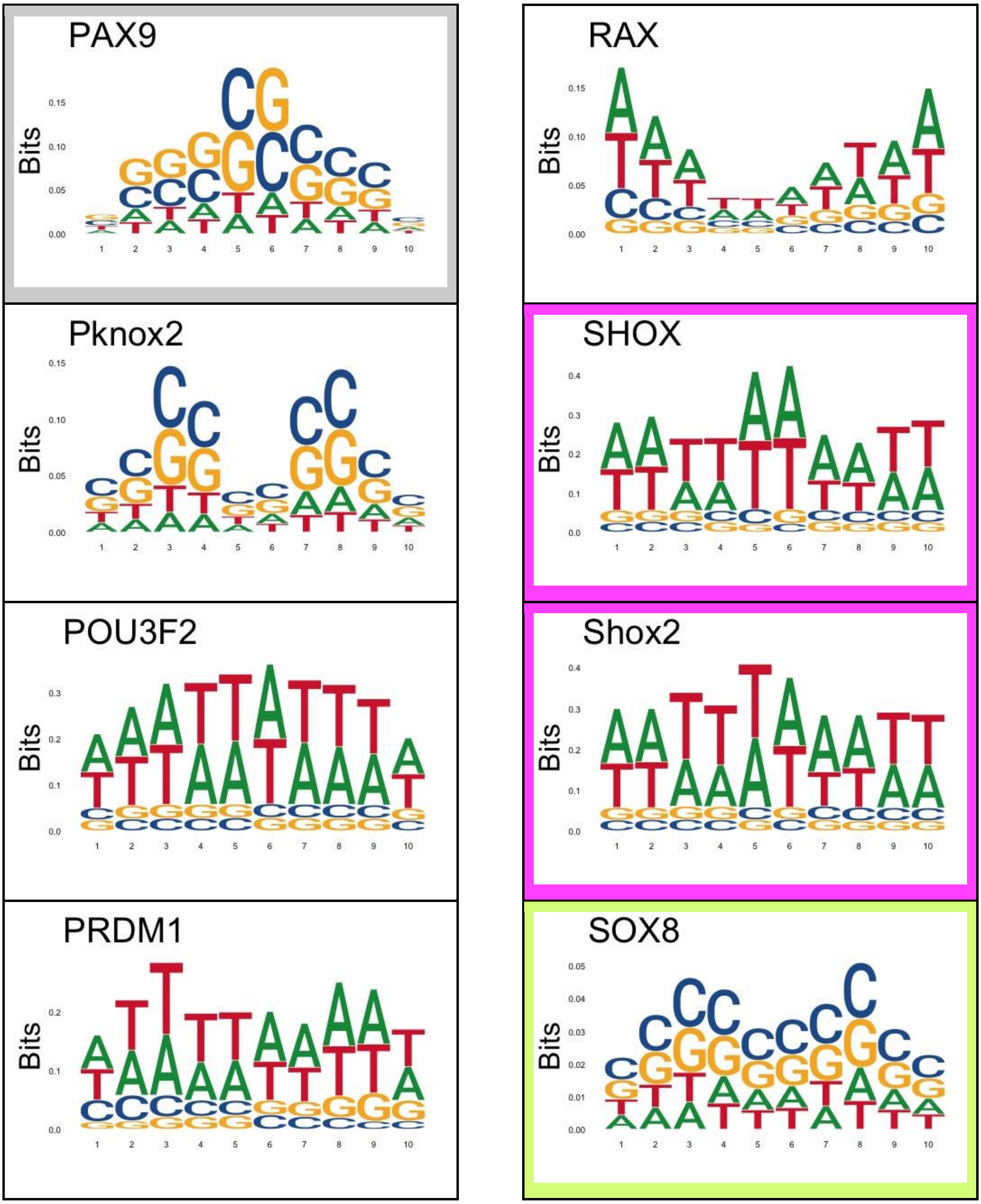
PAX9
RAX
Pknox2
SHOX
POU3F2
Shox2
PRDM1
SOX8
Bits

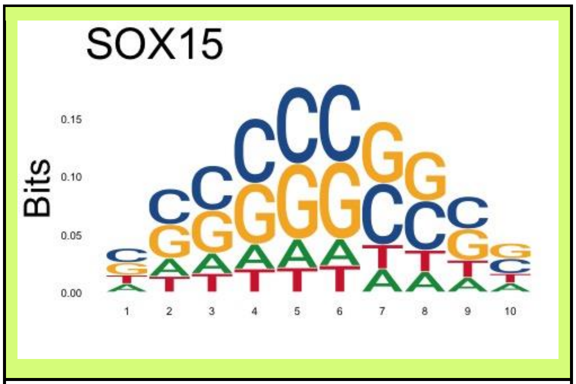
SOX15
Bits
0.15
0.10
0.05
0.00
1 2 3 4 5 6 7 8 9 10

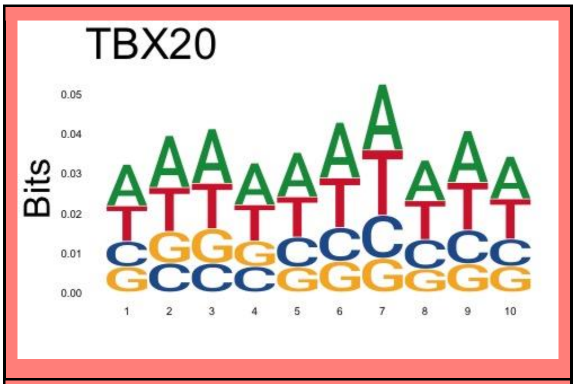
TBX20
Bits
0.05
0.04
0.03
0.02
0.01
0.00
1 2 3 4 5 6 7 8 9 10

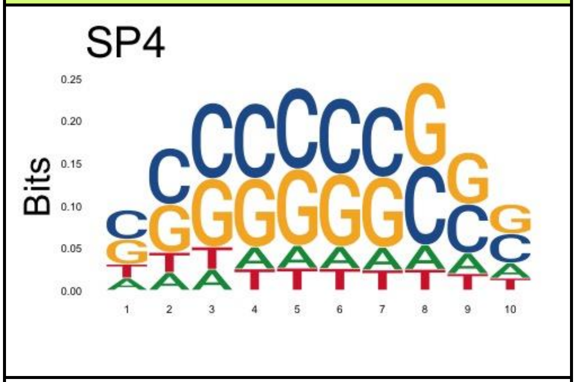
SP4
Bits
0.25
0.20
0.15
0.10
0.05
0.00
1 2 3 4 5 6 7 8 9 10

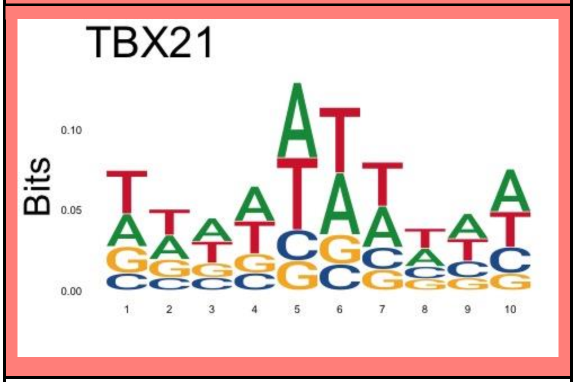
TBX21
Bits
0.10
0.05
0.00
1 2 3 4 5 6 7 8 9 10

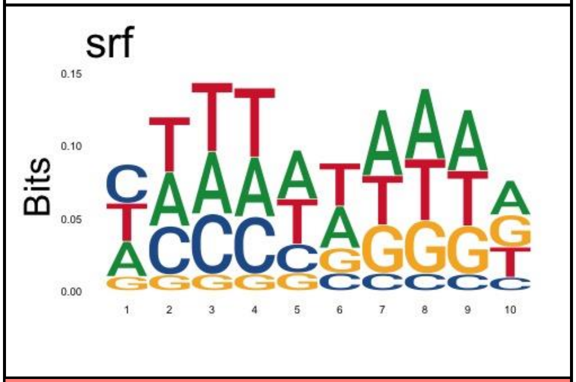
srf
Bits
0.15
0.10
0.05
0.00
1 2 3 4 5 6 7 8 9 10

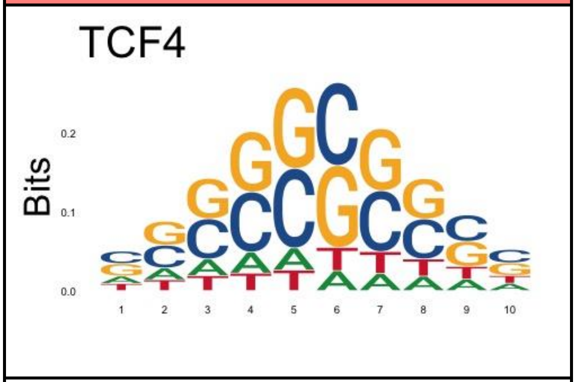
TCF4
Bits
0.2
0.1
0.0
1 2 3 4 5 6 7 8 9 10

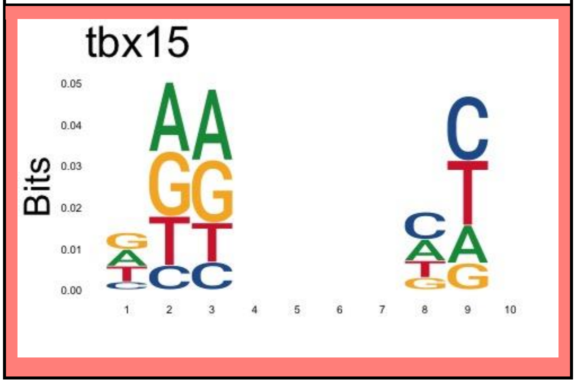
tbx15
Bits
0.05
0.04
0.03
0.02
0.01
0.00
1 2 3 4 5 6 7 8 9 10

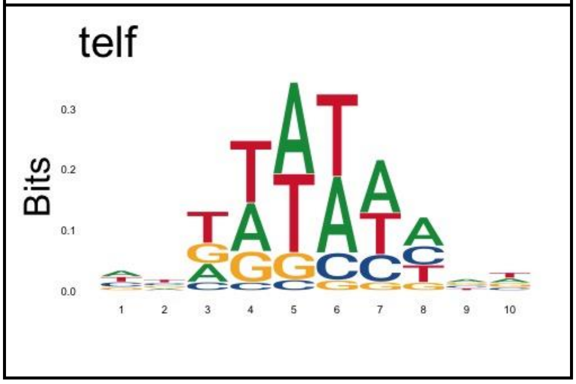
telf
Bits
0.3
0.2
0.1
0.0
1 2 3 4 5 6 7 8 9 10

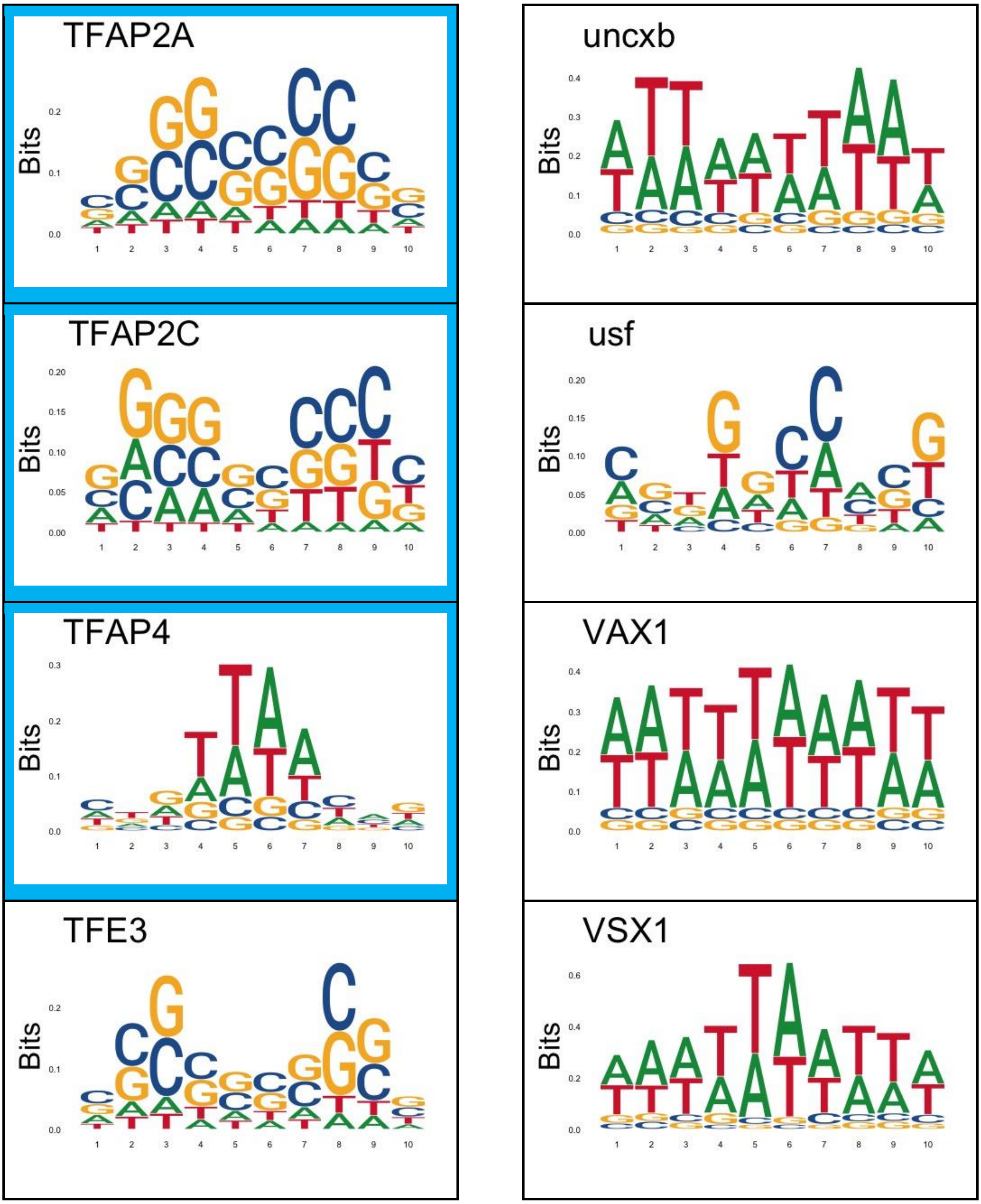
TFAP2A
uncxb
TFAP2C
usf
TFAP4
VAX1
TFE3
VSX1
Bits

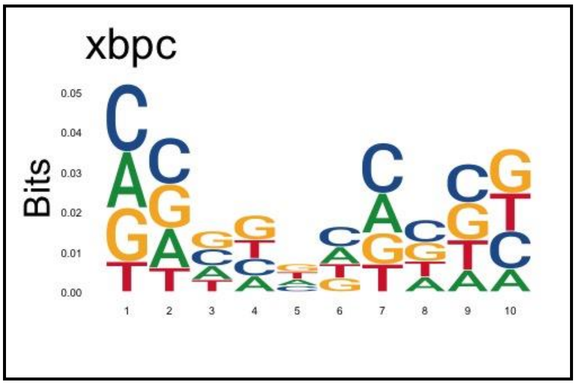


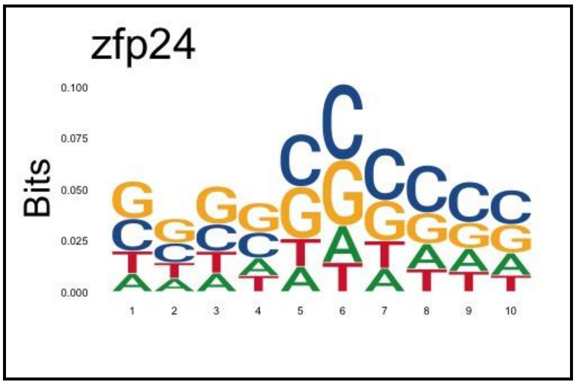


**Supplementary Table S4.** MSE calculated for the results obtained using DNAaffinity, CRPTS/CRPT, DNAShapeR and DeepSELEX using the uPMB and HT-SELEX data respectively.

| Method | DATA | MSE |
|---|---|---|
| DNAffinity | HT-SELEX | 0.0005 ± 0.0002 |
| DEEPSELEX | HT-SELEX | 0.013 ± 0.008 |
| DNAShapeR | HT-SELEX | 0.034 ± 0.029 |
| DNAffinity | uPBM | 0.011 ± 0.004 |
| CRPTS/CRPT | uPBM | 0.16 ± 0.112 |
| DNAShapeR | uPBM | 0.008 ± 0.011 |
